# Solution conformational differences between conventional and CENP-A nucleosomes are accentuated by reversible deformation under high pressure

**DOI:** 10.1101/2025.01.16.633457

**Authors:** Kushol Gupta, Nikolina Sekulić, Praveen Kumar Allu, Nicklas Sapp, Qingqiu Huang, Kathryn Sarachan, Mikkel Christensen, Reidar Lund, Susan Krueger, Joseph E. Curtis, Richard E. Gillilan, Gregory D. Van Duyne, Ben E. Black

## Abstract

Solution-based interrogation of the physical nature of nucleosomes has its roots in X-ray and neutron scattering experiments, including those that provided the initial observation that DNA wraps around core histones. In this study, we performed a comprehensive small-angle scattering study to compare canonical nucleosomes with variant centromeric nucleosomes harboring the histone variant, CENP-A. We used nucleosome core particles (NCPs) assembled on an artificial positioning sequence (Widom 601) and compared these to those assembled on a natural α-satellite DNA cloned from human centromeres. We establish the native solution properties of octameric H3 and CENP-A NCPs using analytical ultracentrifugation (AUC), small-angle X-ray scattering (SAXS), and contrast variation small-angle neutron scattering (CV-SANS). Using high-pressure SAXS (HP-SAXS), we discovered that both histone identity and DNA sequence have an impact on the stability of octameric nucleosomes in solution under high pressure (300 MPa), with evidence of reversible unwrapping in these experimental conditions. Both canonical nucleosomes harboring conventional histone H3 and their centromeric counterparts harboring CENP-A have a substantial increase in their radius of gyration, but this increase is much less prominent for centromeric nucleosomes. More broadly for chromosome-related research, we note that as HP-SAXS methodologies expand in their utility, we anticipate this will provide a powerful solution-based approach to study nucleosomes and higher-order chromatin complexes.

## Introduction

Just as the canonical nucleosome is the fundamental repeating unit of chromatin, the specialized nucleosome harboring the histone H3 variant CENP-A is, in general, the repeating unit of chromatin at the portion of the centromere that serves as the foundation of the mitotic kinetochore (Palmer et al. 1987; Kixmoeller, Allu, and Black 2020; Yatskevich, Barford, and Muir 2023). The roles of the centromere also include serving as the final location of sister chromatin cohesion until anaphase onset, and as the site for enrichment in early mitosis of the chromosome passenger complex where it functions to monitor the presence and quality of spindle microtubule attachments to the kinetochore (Broad and DeLuca 2020; Carmena et al. 2012). Outside of mitosis, CENP-A nucleosomes serve to epigenetically maintain the location of the centromere. Throughout the cell cycle, CENP-A nucleosomes bind to a sixteen-subunit protein complex called the constitutive centromere-associated network (CCAN) (Foltz et al. 2006; Okada et al. 2006). A CENP-A nucleosome with steady-state wrapping of ∼1.7 turns of DNA, similar to conventional nucleosomes, is evident in structural analyses of reconstituted CENP-A nucleosome/CCAN complexes (Yan et al. 2019; Yatskevich et al. 2022) and in the context of their natural counterparts at the centromeres of intact human chromosomes (Kixmoeller, Chang, and Black 2025). That said, solution-based studies have revealed that CENP-A nucleosomes have rigid histone cores (Black et al. 2004; Black et al. 2007; Sekulic et al. 2010) and loose superhelical DNA termini (Hasson et al. 2013; Falk et al. 2015; Conde e Silva et al. 2007). The methodologies that have been used to study CENP-A nucleosomes have been limited in scope, though, and it is likely there are other distinguishing physical features of CENP-A nucleosomes that could impact their essential functions at the centromere.

In this light, it is important to note that high-resolution methods typically provide a static view of atomic structure, hence limiting insights into intrinsic flexibility and ensemble-averaged dynamic properties such as conformational changes and interdomain flexibility. Since the 1970s, small-angle scattering (SAS) has served as an invaluable complementary tool for structural biology that has bridged the gap between atomic-level structure and solution behavior (Weiss 2017; Mohammed, Soloviov, and Jeffries 2024). The angular-dependent decay in scattering intensity provides rich model-independent information about macromolecular size, shape, and flexibility in solution, and with the availability of experimental and *ab initio* atomic structures, a rigorous test of solution structure.

The first studies of nucleosomes using SAS arrived in the 1970s (Olins and Olins 1974; Baldwin et al. 1975; Stuhrmann and Duee 1975; Finch and Klug 1976; Woodcock, Safer, and Stanchfield 1976; Hjelm et al. 1977), when neutron contrast variation (Krueger 2022) experiments were first employed to distinguish between protein and DNA components of the nucleosome. These early studies revealed the fundamental properties of chromatin and the nucleosome, such as the size, shape, and quaternary arrangement. Canonical small-angle X-ray scattering (SAXS) and neutron scattering (SANS) experiments during this era contributed to the initial development of the solenoidal (Finch and Klug 1976) and “beads-on-a-string” (Olins and Olins 1974; Woodcock, Safer, and Stanchfield 1976) models of chromatin structure. These studies also established the now well-known structural feature of DNA wrapping around an inner histone octamer in mononucleosomes (Hjelm et al. 1977). Advancements in deuterium labeling techniques in the mid -1970s and 1980s enabled subsequent higher-resolution SANS studies, offering deeper insights into nucleosome assembly and component interactions (Baldwin et al. 1975; Moore 1982). Similarly, the advent of synchrotron SAXS in the 1990s improved signal-to-noise ratios, enabling more detailed investigations into chromatin flexibility and conformational changes in response to factors such as salt concentration (Hansen et al. 2017; Nishino et al. 2012; Maeshima et al. 2016; Joti et al. 2012). Over the past two decades, integrative approaches combining SAXS with high-resolution structural modeling methods have driven further advances in understanding. These approaches have allowed detailed investigations into the effects of histone variants (Tachiwana et al. 2011; Sugiyama et al. 2015), nucleosome interactors (Yang et al. 2011), post-translational modifications (PTMs) (Brehove et al. 2015), and DNA sequence (Yang et al. 2011) on nucleosome structure. X-ray scattering has also shed light on nucleosome dynamics, remodeling processes, and the organization of higher-order chromatin assemblies. However, despite the early successes of SANS with contrast variation (CV-SANS) in nucleosome studies, relatively few such studies have been published since over the past 30 years. This is surprising given significant advancements in neutron reactor sources, detector technology, data analysis techniques, and the availability of high-resolution atomic structures through this time (Ashkar et al. 2018). While SANS has found increasing applications in the biological realm (Ashkar et al. 2018), most contrast variation studies performed during this period have focused on other large assemblies or specific protein-DNA interactions, leaving nucleosome-specific applications still relatively underexplored.

In this study, we revisit these canonical applications of SAS, including contrast variation SANS (CV-SANS), and introduce the emerging state-of-the-art approach of high-pressure small-angle X-ray scattering (HP-SAXS) to investigate the structural properties of nucleosome core particles (NCPs), with a focus on centromeric chromatin. We examine how differences in protein composition and DNA sequence influence NCP structure and dynamics in solution. Specifically, we compare canonical H3.1 histone octamers with centromeric histone octamers, in which the canonical H3.1 is replaced by the variant centromeric histone CENP-A (approximately ∼63% sequence identity in the histone core but highly divergent histone tails). These octamers are assembled on two different and well-studied DNA sequences: the strong-positioning Widom DNA (601) sequence (Lowary and Widom 1998) and an AT-rich α-satellite sequence derived from centromeric DNA(Harp et al. 1996). Using SAXS and SANS, we demonstrate that NCPs containing CENP-A and α-satellite DNA exhibit a measurably larger spatial extent and evidence of flexibility compared to canonical H3 NCPs, in agreement with prior studies. Analytical ultracentrifugation (AUC) studies of α-satellite NCPs confirm their octameric stoichiometry and reveal physical polydispersity specific to the combination of CENP-A histones and α-satellite DNA. To further probe these differences, we employ HP-SAXS and discover a reversible sensitivity of α-satellite NCPs to pressure, highlighting structural and dynamic properties specific to nucleosome composition.

## Results

### Small-angle X-ray scattering (SAXS) reveals differences in the spatial extent of nucleosomes assembled with different histone and DNA sequences

For our SAXS experiments, we reconstituted the canonical H3.1 and CENP-A-derived histone octamers on either the well-studied strong positioning Widom 601 sequence (Lowary and Widom 1998) (H3-601 and CENP-A-601, respectively) or on native α-satellite DNA (Harp et al. 1996) (H3-αSat or CENP-A-αSat, respectively) to determine their structural properties in solution. The strong positioning on Widom 601 is thought to be due to dinucleotide pair combinations that accommodate the deformations in DNA path conferred by histone DNA wrapping (Lowary and Widom 1998). It is reasonable to expect that while the overall path of DNA wrapping will be largely similar in Widom 601 and natural DNA sequences (Allu et al. 2019; Vasudevan, Chua, and Davey 2010; Wang, Xiong, and Cramer 2021), there could nonetheless be DNA sequence-dependent deviations in local structure and dynamics. We first probed the structural properties of these reconstituted NCPs in solution using small-angle X-ray scattering (SAXS), a technique that is very sensitive to changes in macromolecular conformation in solution. Preliminary measurements were performed on a rotating anode X-ray source to optimize buffer and concentration ranges and to eliminate any effects of interparticle interference (see **Methods**). Final synchrotron data were recorded at multiple concentrations in the range of 0.5 mg/mL - 1.2 mg/mL. Classical Guinier analyses indicated particles free of self-association or aggregation (**Figure 1A**), and together these measurements yielded structural parameters from SAXS consistent with prior literature reports on canonical particles (Yang et al. 2011; Hjelm et al. 1977; Sugiyama et al. 2014; Sugiyama et al. 2015) (**Table 1**). We find that H3 and CENP-A NCPs assembled on 601 sequences have similar radii of gyration (R_g_ ∼41 Å-42 Å). An indirect Fourier transformation of the primary data into shape distribution functions (P(r)) allows for a direct comparison of the distribution of interatomic vectors within the scattering volumes in real-space. While both 601 particles show a very similar P(r) profiles, difference analysis (**Figure 1B**) reveals a shift in interatomic vectors away from the middle of the distribution when the CENP-A-601 particle is compared to the H3-601 particle. The larger maximum dimensions (D_max_) determined (∼122 Å-140 Å) for CENP-A-601 coincides with a decrease in interatomic vectors between ∼60 Å-100 Å and the overall slight increase in R_g_ when compared to H3-601 (D_max_ ∼118 Å-125 Å) (**Table 1**). On α-satellite DNA, the R_g_ of H3 on α-satellite DNA is more like those observed for 601 NCPs. In contrast, CENP-A nucleosomes on α-satellite DNA show significantly larger spatial extents (R_g_ = ∼43 Å - 44 Å). Interestingly, both the H3 and CENP-A α-satellite NCPs have larger maximum dimensions (∼126 and 155 Å, respectively) with CENP-A showing the largest interatomic distances observed.

**Figure 1.**
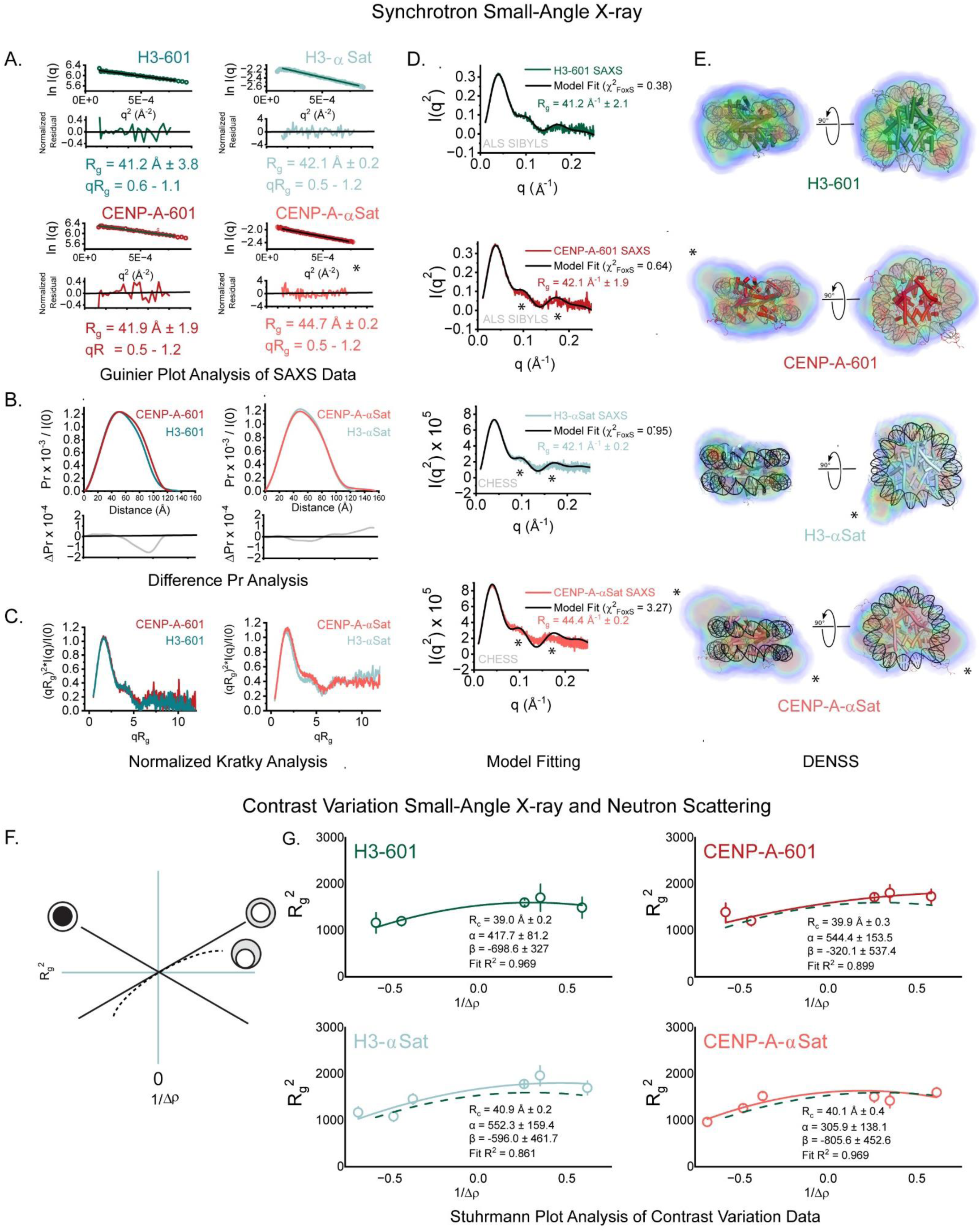
SAXS and SANS measurements reveal subtle dynamic changes between NCPs conferred by histone and DNA composition. **A. Guinier Plots.** Guinier plot analyses (lnI(*q*) vs. *q*^2^) of SAXS data (dots) for NCPs, with residuals from the fitted lines shown below [colors]. Monodispersity is evidenced by linearity in the Guinier region of the scattering data and agreement of the I_0_ and R_g_ values determined with inverse Fourier transform analysis by the programs GNOM (**Table 1**). Guinier analyses were performed where qR_g_ ≤ 1.2. **B. Difference P(r). C. Normalized Kratky Plots. D. FoxS Fitting.** The recorded X-ray intensity for H3-601 (green line) is shown as a function of *q* (*q*=4πsinθ/λ, where 2θ is the scattering angle) on a Kratky plot (I*q*^2^ vs *q*), to emphasize the features in the middle *q* regime associated with conformational changes in solution. Parameters derived from SAXS analyses are summarized in **Table 1**. Show as a solid black line is the fit calculated intensity profile from the H3-601 atomistic model obtained using the program FoxS^65^, with the χ^2^ associated with the fitting provided. Similar analyses are provided in panels **C-E** for CENP-A-601 (C, light red, versus the CENP-A-601 model), H3-αSat (D, cyan, versus the H3-601 model), and CENP-A-αSat (E, cyan, versus the CENP-A-601 model). Relative to the H3-601 model fit, the other three particles examined show differing levels of discrepancy at the ∼0.1 and ∼0.16 *q* peak features, and larger R_g_ values, indicating modest differences in solution conformation. **E. DENSS analysis of synchrotron SAXS data.** Shown in orthogonal views for each of the four particles examined by SAXS are *ab initio* electron density reconstructions, docked with the corresponding H3 or CENP-A nucleosome model. Arrows denote spatial discrepancies between the experimental volumes and idealized models that correlate strongly to changes in DNA end positioning. **F**. **Schematic of Stuhrmann Plot contrast variation profiles for different idealized structures**. In each case, higher scattering density in the composite particle is indicated by darker shading. The x-intercept at zero provides the R_g_ at infinite contrast (R_c_). In profiles where the slope is positive (α>0), the higher density component is located on the periphery of the composite particle, whereas negative slopes (α<0) indicated the opposite. For non-linear profiles (where β≠0), the two components are askew relative to the center of mass, as illustrated. **G**. **Stuhrmann Plot analysis (R_g_^2^ vs Δρ^−1^)** for the H3-601 particle is shown (dark red), with four SANS data points and one SAXS data point fit with the Stuhrmann equation. Shown for each data point are the errors associated with classical Guinier fitting (see Table 2). Provided in the panel are the numerical parameters derived from the fitting of the Stuhrmann equation, including the R^2^ from the fitting. Similar analyses are provided in panels for CA-601 (I, green, four SANS data points and one SAXS data point), H3-αSat (J, cyan, five SANS data points and one SAXS data point), and CA-αSat (K, light red, five SANS data points and one SAXS data point), with the H3-601 Stuhrmann result shown as a dotted dark red line for comparison. Relative to H3-601, the other three particles have slightly increased spatial extent, as indicated by a modest upwards translation along the y-axis. All four particles have very similar determined R_c_ values, and positive α and β terms, indicate very similar gross compositional distributions of the protein and DNA components in these experimental conditions. All numerical parameters derived from these analyses and additional supporting measures are also provided in **Supplemental Figure 1** and **Supplemental Tables 2** and **Supplemental Table 3**.

**Table 1:**
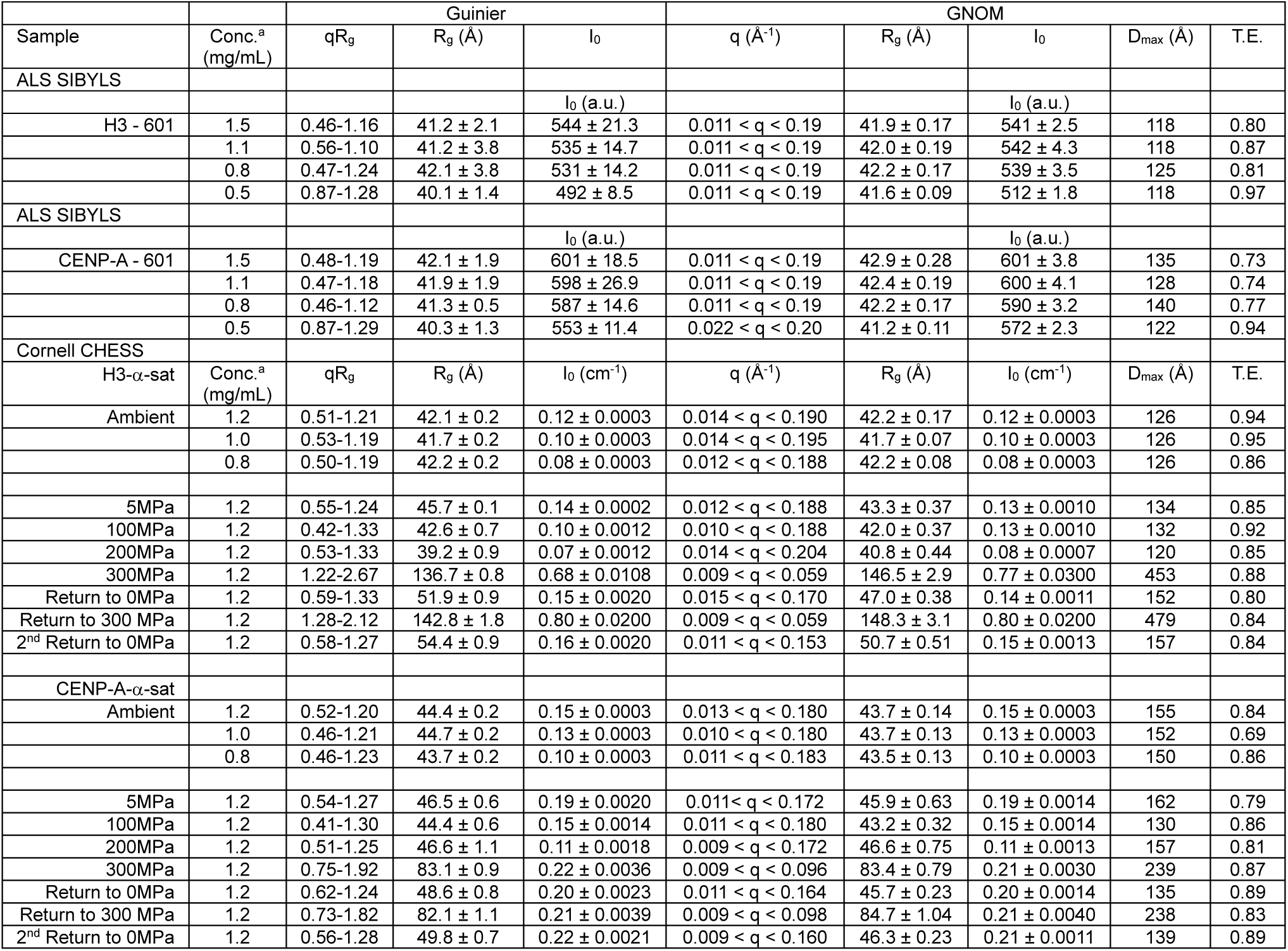
Table of Extended Parameters derived from Small-angle X-ray Scattering (SAXS) Analysis.

| Sample | Conc.<sup>a</sup><br>(mg/mL) | Guinier |  |  | GNOM |  |  |  |  |
| --- | --- | --- | --- | --- | --- | --- | --- | --- | --- |
|  |  | qR<sub>g</sub> | R<sub>g</sub> (Å) | I<sub>0</sub> | q (Å<sup>-1</sup>) | R<sub>g</sub> (Å) | I<sub>0</sub> | D<sub>max</sub> (Å) | T.E. |
| ALS SIBYLS |  |  |  |  |  |  |  |  |  |
|  |  |  |  | I<sub>0</sub> (a.u.) |  |  | I<sub>0</sub> (a.u.) |  |  |
| H3 - 601 | 1.5 | 0.46-1.16 | 41.2 ± 2.1 | 544 ± 21.3 | 0.011 < q < 0.19 | 41.9 ± 0.17 | 541 ± 2.5 | 118 | 0.80 |
|  | 1.1 | 0.56-1.10 | 41.2 ± 3.8 | 535 ± 14.7 | 0.011 < q < 0.19 | 42.0 ± 0.19 | 542 ± 4.3 | 118 | 0.87 |
|  | 0.8 | 0.47-1.24 | 42.1 ± 3.8 | 531 ± 14.2 | 0.011 < q < 0.19 | 42.2 ± 0.17 | 539 ± 3.5 | 125 | 0.81 |
|  | 0.5 | 0.87-1.28 | 40.1 ± 1.4 | 492 ± 8.5 | 0.011 < q < 0.19 | 41.6 ± 0.09 | 512 ± 1.8 | 118 | 0.97 |
| ALS SIBYLS |  |  |  |  |  |  |  |  |  |
|  |  |  |  | I<sub>0</sub> (a.u.) |  |  | I<sub>0</sub> (a.u.) |  |  |
| CENP-A - 601 | 1.5 | 0.48-1.19 | 42.1 ± 1.9 | 601 ± 18.5 | 0.011 < q < 0.19 | 42.9 ± 0.28 | 601 ± 3.8 | 135 | 0.73 |
|  | 1.1 | 0.47-1.18 | 41.9 ± 1.9 | 598 ± 26.9 | 0.011 < q < 0.19 | 42.4 ± 0.19 | 600 ± 4.1 | 128 | 0.74 |
|  | 0.8 | 0.46-1.12 | 41.3 ± 0.5 | 587 ± 14.6 | 0.011 < q < 0.19 | 42.2 ± 0.17 | 590 ± 3.2 | 140 | 0.77 |
|  | 0.5 | 0.87-1.29 | 40.3 ± 1.3 | 553 ± 11.4 | 0.022 < q < 0.20 | 41.2 ± 0.11 | 572 ± 2.3 | 122 | 0.94 |
| Cornell CHESS |  |  |  |  |  |  |  |  |  |
| H3-α-sat | Conc.<sup>a</sup><br>(mg/mL) | qR<sub>g</sub> | R<sub>g</sub> (Å) | I<sub>0</sub> (cm<sup>-1</sup>) | q (Å<sup>-1</sup>) | R<sub>g</sub> (Å) | I<sub>0</sub> (cm<sup>-1</sup>) | D<sub>max</sub> (Å) | T.E. |
| Ambient | 1.2 | 0.51-1.21 | 42.1 ± 0.2 | 0.12 ± 0.0003 | 0.014 < q < 0.190 | 42.2 ± 0.17 | 0.12 ± 0.0003 | 126 | 0.94 |
|  | 1.0 | 0.53-1.19 | 41.7 ± 0.2 | 0.10 ± 0.0003 | 0.014 < q < 0.195 | 41.7 ± 0.07 | 0.10 ± 0.0003 | 126 | 0.95 |
|  | 0.8 | 0.50-1.19 | 42.2 ± 0.2 | 0.08 ± 0.0003 | 0.012 < q < 0.188 | 42.2 ± 0.08 | 0.08 ± 0.0003 | 126 | 0.86 |
| 5MPa | 1.2 | 0.55-1.24 | 45.7 ± 0.1 | 0.14 ± 0.0002 | 0.012 < q < 0.188 | 43.3 ± 0.37 | 0.13 ± 0.0010 | 134 | 0.85 |
| 100MPa | 1.2 | 0.42-1.33 | 42.6 ± 0.7 | 0.10 ± 0.0012 | 0.010 < q < 0.188 | 42.0 ± 0.37 | 0.13 ± 0.0010 | 132 | 0.92 |
| 200MPa | 1.2 | 0.53-1.33 | 39.2 ± 0.9 | 0.07 ± 0.0012 | 0.014 < q < 0.204 | 40.8 ± 0.44 | 0.08 ± 0.0007 | 120 | 0.85 |
| 300MPa | 1.2 | 1.22-2.67 | 136.7 ± 0.8 | 0.68 ± 0.0108 | 0.009 < q < 0.059 | 146.5 ± 2.9 | 0.77 ± 0.0300 | 453 | 0.88 |
| Return to 0MPa | 1.2 | 0.59-1.33 | 51.9 ± 0.9 | 0.15 ± 0.0020 | 0.015 < q < 0.170 | 47.0 ± 0.38 | 0.14 ± 0.0011 | 152 | 0.80 |
| Return to 300 MPa | 1.2 | 1.28-2.12 | 142.8 ± 1.8 | 0.80 ± 0.0200 | 0.009 < q < 0.059 | 148.3 ± 3.1 | 0.80 ± 0.0200 | 479 | 0.84 |
| 2<sup>nd</sup> Return to 0MPa | 1.2 | 0.58-1.27 | 54.4 ± 0.9 | 0.16 ± 0.0020 | 0.011 < q < 0.153 | 50.7 ± 0.51 | 0.15 ± 0.0013 | 157 | 0.84 |
| CENP-A-α-sat | | | | | | | | | |
| Ambient | 1.2 | 0.52-1.20 | 44.4 ± 0.2 | 0.15 ± 0.0003 | 0.013 < q < 0.180 | 43.7 ± 0.14 | 0.15 ± 0.0003 | 155 | 0.84 |
|  | 1.0 | 0.46-1.21 | 44.7 ± 0.2 | 0.13 ± 0.0003 | 0.010 < q < 0.180 | 43.7 ± 0.13 | 0.13 ± 0.0003 | 152 | 0.69 |
|  | 0.8 | 0.46-1.23 | 43.7 ± 0.2 | 0.10 ± 0.0003 | 0.011 < q < 0.183 | 43.5 ± 0.13 | 0.10 ± 0.0003 | 150 | 0.86 |
| 5MPa | 1.2 | 0.54-1.27 | 46.5 ± 0.6 | 0.19 ± 0.0020 | 0.011 < q < 0.172 | 45.9 ± 0.63 | 0.19 ± 0.0014 | 162 | 0.79 |
| 100MPa | 1.2 | 0.41-1.30 | 44.4 ± 0.6 | 0.15 ± 0.0014 | 0.011 < q < 0.180 | 43.2 ± 0.32 | 0.15 ± 0.0014 | 130 | 0.86 |
| 200MPa | 1.2 | 0.51-1.25 | 46.6 ± 1.1 | 0.11 ± 0.0018 | 0.009 < q < 0.172 | 46.6 ± 0.75 | 0.11 ± 0.0013 | 157 | 0.81 |
| 300MPa | 1.2 | 0.75-1.92 | 83.1 ± 0.9 | 0.22 ± 0.0036 | 0.009 < q < 0.096 | 83.4 ± 0.79 | 0.21 ± 0.0030 | 239 | 0.87 |
| Return to 0MPa | 1.2 | 0.62-1.24 | 48.6 ± 0.8 | 0.20 ± 0.0023 | 0.011 < q < 0.164 | 45.7 ± 0.23 | 0.20 ± 0.0014 | 135 | 0.89 |
| Return to 300 MPa | 1.2 | 0.73-1.82 | 82.1 ± 1.1 | 0.21 ± 0.0039 | 0.009 < q < 0.098 | 84.7 ± 1.04 | 0.21 ± 0.0040 | 238 | 0.83 |
| 2<sup>nd</sup> Return to 0MPa | 1.2 | 0.56-1.28 | 49.8 ± 0.7 | 0.22 ± 0.0021 | 0.009 < q < 0.160 | 46.3 ± 0.23 | 0.21 ± 0.0011 | 139 | 0.89 |

**Table 2.**
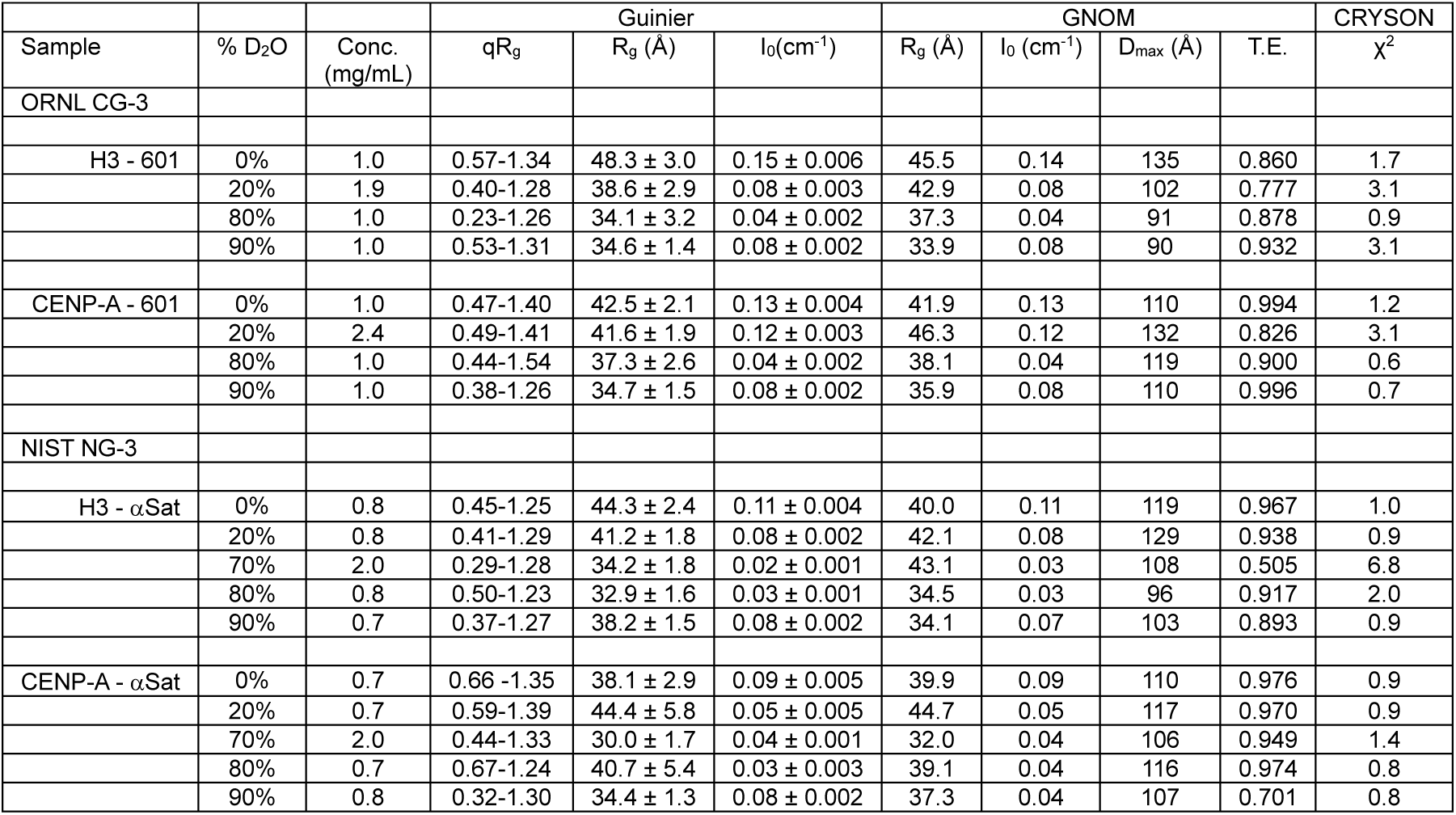
Parameters derived from Small-angle Neutron Scattering (SANS) analysis.

Normalized Kratky representations of the primary data ((*qR_g_*)^2^ *x I(q)/I_0_) vs qR_g_*) (Receveur-Brechot and Durand 2012) were employed to facilitate direct comparison in a model-independent fashion, placing emphasis on the length scales in the middle *q* regime (∼ 0.05 Å^−1^ < *q* < 0.2 Å^−1^) that are most strongly correlated to changes in macromolecular conformation, and for overall insight into the compactness and overall shape. The normalized Kratky profiles for H3-601 and CENP-A-601 are shown in **Figure 1C**. In both cases, a lower primary peak feature is observed where *qR_g_* ∼1.7 (Putnam et al. 2007; Receveur-Brechot and Durand 2012), consistent with globular/compact particles and a properly determined R_g_ from Guinier analysis, followed by a second smaller peak feature at ∼4 *qR_g_* before a return to a baseline intensity, all indicating a well-folded and compact macromolecule. In this comparison, only modest differences can be discerned in the middle *q* regime in a pairwise comparison. In contrast, after the lower primary peak feature at ∼1.7 *qR_g_*, the profiles for α-satellite DNA NCPs vary upward as a function of *qR_g_* with no return to baseline at higher *q* values, consistent with the increased R_g_ and D_max_ values observed and indicative of a more dynamic behavior in solution associated with larger particle volumes (Rambo and Tainer 2011).

To more precisely relate the observed solution properties obtained from these preparations to canonical atomic structures, we also generated all-atom models of both H3-601 and CENP-A-601 using molecular dynamics. Our H3-601 model is derived from the PDB 3LZ0 crystal structure (Kato et al. 2011), with unresolved histone tails (comprising about ∼30% of the total protein mass in a nucleosome core particle) modeled as collapsed random coils (see **Methods**). Similarly, the CENP-A model was constructed using the PDB 3AN2 atomic structure(Tachiwana et al. 2011) as a template for the protein component. Full atomic coordinates for the models including histone tails and correct DNA sequence were built manually and models were minimized (see Material and Methods) To compare our experimental scattering results to these models, we employed the Debye relationship (Debye 1915). However, the proper interpretation of experimental intensity profiles and their reconciliation with atomistic models of composite particles like protein-DNA complexes requires accurate predictions of the hydration layers and excluded solvent (Knight and Hub 2015; Svergun, Barberato, and Koch 1995a) due to differences in the scattering length densities (SLDs) between protein and DNA in X-rays, as well as nonuniform hydration layers (Poitevin et al. 2011) which contribute to the overall X-ray scattering. To address these considerations, the program WAXSiS (Knight and Hub 2015) was employed to explicitly calculate the hydration layer via molecular dynamics simulation. These fits were compared to calculations performed with a more common implicit solvent model where the boundary layer is an adjustable parameter in the fitting (FoxS (Schneidman-Duhovny et al. 2016)). Using both algorithms, general agreement was observed between our atomistic models on 601 DNA and the corresponding solution scattering from 601 and α-satellite DNA containing particles, using standard Kratky plots (*Iq^2^ vs q*) to again place focus on middle *q* features (**Figure 1D and Supplemental Figure 1**). The H3-601 particle was concordant with a canonical atomic model across scattering angles where q_max_<0.3 Å^−1^, whereas with the CENP-A-601 particle, only fine discrepancies were readily apparent, mapping primarily to peak features at ∼0.1 *q* (corresponding to a length scale of ∼63 Å, where d = *2π/q*,). In contrast, direct comparison of our respective canonical models to α-satellite NCP data show less concordance in middle q. While the H3-αSat particle fit is somewhat concordant (χ^2^=0.95) with modest discrepancy most apparent near the ∼0.16 *q* peak feature (corresponding to a length scale of ∼39.3 Å), CENP-A-α-Sat data shows the poorest correlations overall, with the CENP-A-601 atomic model poorly matching across nearly all of the middle *q* regime (χ^2^=3.27).

To further visualize the changes in canonical structure detected, *ab initio* modeling approaches were applied using the SAXS data to generate low-resolution particle envelopes of the solution average ensemble. The algorithm DENSS (Grant 2018b) allows for the reconstruction of the electron density of particles at low resolution, allowing for a real space assessment of available atomic models. We applied this algorithm to data from the four particles and the results are shown in **Figure 1E**. The molecular envelopes were generated with no symmetry restraints, as to not bias the shapes derived (see **Methods**). Consistent with direct fitting of the experimental profiles, strong spatial correlation was observed for the H3-601 particle when the structure is docked into the calculated volume. The strongest density contours in this reconstruction correlates well with the DNA component of the models, consistent with the dominance of the DNA signal in the X-ray measurements. Relative to this result, the DENSS calculation for the CENP-A-601 particle yielded a particle shape with greater oblate character and more apparent asymmetry. Docking of the CENP-A-601 model into this SAXS-derived volume is suggestive of differences in the DNA ends of the particle, consistent with prior reports of DNA entry/exit behavior (Conde e Silva et al. 2007), greater micrococcal nuclease sensitivity of these regions (Bloom and Carbon 1982; Falk et al. 2015; Hasson et al. 2013) and the atomic structure of CENP-A-601 (Tachiwana et al. 2011; Ali-Ahmad et al. 2019) where these regions are entirely disordered and unresolved in the electron density. *Ab initio* reconstructions of H3-αSat show reasonable spatial correlation with the H3-601 canonical model, and like the CENP-A-601 result, indicate asymmetry corresponding to the positioning of DNA ends when docked. The CENP-A-αSat shows the greatest spatial discrepancy versus a docked canonical model, consistent with significant displacement of both DNA ends. Together, these data indicate that SAXS on NCPs in solution captures dynamic features resulting from the unique combination of histone and DNA sequences reflecting quaternary arrangements that do not coincide well with canonical models. Differences between CENP-A and H3-containing nucleosomes are consistent with flexible DNA ends previously observed by micrococcal nuclease digestion, X-ray crystallography, and cryo-electron microscopy, and are most pronounced in the CENP-A derived species.

### Contrast Variation Small-Angle Neutron Scattering (CV-SANS) measures the gross compositional distribution of NCPs

A strength of the SAXS approach in studying NCPs is the higher scattering power of the DNA component in X-rays and higher signal-to-noise across larger scattering angles, making it very well suited to detect differences in DNA conformation. While it is possible to entirely contrast away the protein contribution of composite particles in X-rays using excipients such as glycerol or sucrose (Chen et al. 2014; Mauney et al. 2021; Mauney et al. 2018), such approaches limit insights attained by contrast variation to the DNA component only, as it exists in the complex via a relatively narrow window for contrast variation. Hence, SAXS alone is limited in its ability to discern the properties of the protein component of these assemblies. Contrast variation studies using small-angle neutron scattering (CV-SANS) provide a powerful complement to the X-ray approach and can provide a wider range of accessible contrast using specific mixtures of H_2_O and D_2_O. Due to the negative scattering density of H_2_O (−0.56 x 10^−6^ Å^−2^) compared to the positive scattering density of its isotope D_2_O (6.67x 10^−6^ Å^−2^), it is possible to adjust the neutron scattering of an aqueous solution by mixing H_2_O and D_2_O in different ratios so that the scattering of the solution completely coincides with that of the protein (∼2-2.5 x 10^−6^ Å^−2^) or DNA component (∼1.8-2.0 x 10^−6^ Å^−2^). If the solution matches the neutron scattering density of the protein, all the scattering collected will come from the DNA component and vice versa. While typically weaker in signal-to-noise for particles of the size of a nucleosome, this approach is sensitive to both composition and spatial extent and provides a direct determination of the distribution of component parts, as they relate to each other in the larger assembly. In contrast to the measurements initially made in the 1970s, more modern technology provided the opportunity to better capture intensity profiles at contrast points normally afflicted with strong incoherent scattering, thus increasing signal-to-noise and allowing a wider accessible *q* range.

Neutron scattering data were recorded at four to five different contrast points (Δρ) for each of the four particles (Table 2). Despite the implementation of more modern technology, attempts to capture data at calculated protein or DNA-only contrasts points were limited by strong incoherent scatter at the low sample concentrations used. Using the recorded intensities at zero angle (I_0_) by SANS, calculated masses (Kuzmanovic et al. 2003) for each of the four NCPs, all samples were consistent with octameric preparations, and experimental total match points were readily determined (**Supplemental Figure 2** and **Supplemental Table 1**). Monodispersity was again confirmed by classical Guinier analyses (**Table 2** and **Supplemental Figures 3&4**). Using both the available SAXS and SANS data combined, the dependence of R_g_ with the contrast of the individual components and their relative positions within the composite particle was determined using classical Stuhrmann analysis:

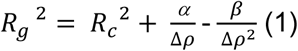

where R_c_ is the radius of gyration (R_g_) at infinite contrast (**Figure 1F**). In the Stuhrmann plots derived for all four NCPs, hyperbolic relationships were apparent, within the error of the determination (**Figure 1G and Supplemental Table 2**), generally consistent with canonical measurements made in 1977 and more recent reports for H3-derived nucleosomes (Hjelm et al. 1977; Sugiyama et al. 2014; Sugiyama et al. 2015). The positive term for α correlates with known structure where the DNA (the denser component) lies at the periphery of the complex, distal from the center-of-mass (Hjelm et al. 1977). The large terms for β describes the relative distribution of scattering length densities within the particle. Distinguishing more recent SANS studies and our studies from the original measurements from 1977 is the ability to model the β term, yielding hyperbolic rather than linear fits in the Stuhrmann plots. In all four cases, similar R_c_ values (R_g_ at infinite contrast) are derived, indicating that within the error associated with this approach, all four NCPs assume a similar spatial extent and gross quaternary structure in these solution conditions. The determined α parameter, which describes the distribution of scattering densities relative to the center of mass, is the smallest for the CENP-A-α-Sat particle, suggesting the greatest changes in the denser DNA component. The term β derived from this fitting relates to the separation of the mass centers of the two components. In our analysis, the determined values are the largest for the CENP-A-αSat, suggesting a more askew position of the protein and DNA components. These results are consistent with the more flexible DNA in CENP-A-αSat sample.

Independent of atomic models, we were readily able to reconcile these contrast variation datasets with empirical core-shell cylinder models using global fitting methods as in employed in the program SASVIEW (Doucet 2017) (**Supplemental Figures 5-6** and **Supplemental Table 4**). Using the experimentally determined SLDs and fixing the radius of the DNA wrap, global fits to the SAXS and SANS data together to this model provided a determination of cylinder radius and length. In this analysis, similar radii (18.1 Å to 19.5 Å) were obtained. Notably, the fit length of the cylinder for the canonical H3-601 particle (59.5 Å ± 2.8 Å) was markedly smaller than those determined for the other three particles (ranging from 68.6 Å ± 0.3 Å to 74.3 Å ± 0.2 Å). The length of the cylinder in this model corresponds well with the nucleosome gyre, which is the path of the DNA as it wraps around the histone protein core, on its smallest dimension. The increase in this fit parameter suggests that the particle gyre is wider in the other three particles, again suggesting a less compact particle in solution. Like with the SAXS data, it is also possible to directly test atomic models against SANS data using the Debye relationship (as implemented in the program CRYSON (Svergun, Barberato, and Koch 1995b)), free of the consideration of solvent boundaries, and with the experimentally determined SLDs from our contrast variation data (**Supplemental Figures 5-6** and **Table 2**). Direct fitting reveals general concordance for most of the experimental data recorded, with the greatest discrepancies observed at the 20% D_2_O (dominated by DNA) and the 70% D_2_O contrast points (dominated by protein, and which were most difficult to capture and fit due to high incoherent scattering). While these results reaffirm composition distribution and the general consistency of canonical structural models to the solution scattering profiles, the SANS data lacks the resolution of our SAXS measurements needed to discern finer changes in DNA conformation, illustrating the complementarity of the two approaches.

### Sedimentation velocity experiments indicate higher physical polydispersity of CENP-A containing nucleosomes

Since α-satellite DNA occurs naturally and harbors both H3 and CENP-A in chromatin, we further focused our investigation on the differences between H3 and CENP-A nucleosomes in the context of α-satellite DNA while H3-601 was used as a reference where needed. To further explore biophysical differences between CENP-A and canonical nucleosomes and to confirm stoichiometry, mass and shape using an orthogonal approach we employed different modalities of analytical ultracentrifugation. To examine the possibility of dissociation of these nucleosomes into hexosomes (Kato et al. 2017), tetrasomes (Sollner-Webb, Camerini-Otero, and Felsenfeld 1976), or other lower-order species, we performed sedimentation equilibrium analysis (SE-AUC), which can provide very precise determinations of molecular weight, independent of any shape effects. Using a mass-averaged partial buoyant density (*ν_bar_*) from known composition and global fitting across multiple rotor speeds and concentrations with strict mass conservation, the masses derived closely matched those expected for DNA-wrapped histone octamers (**Figure 2A** and **Supplemental Tables 4&5**). At these low rotor speeds, both α-Sat particles had determined buoyant masses consistent with octameric NCPs, confirming that the differences observed by SAXS do not correlate to differences in mass and stoichiometry. In agreement with these results and our SAXS analysis, dynamic light scattering (DLS) measurements of the α-Sat particles yielded particle diameters derived from the application of the Stokes-Einstein equation (Einstein 1905) that were generally consistent with the known structure of canonical nucleosome particles and indicative of preparations of high monodispersity (**Figure 2B**).

**Figure 2.**
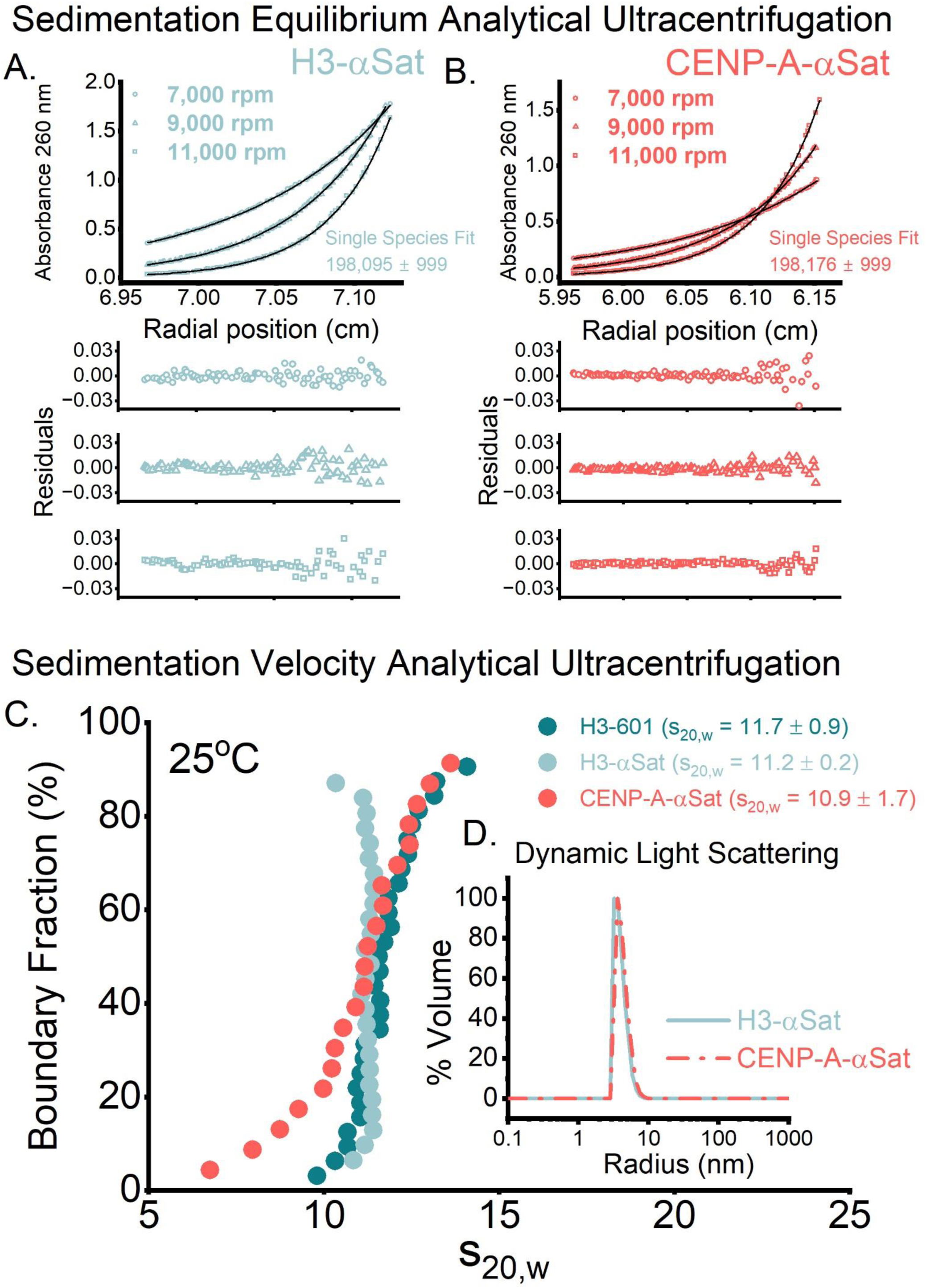
Solution properties of nucleosomes assembled on natural α-satellite DNA probed by analytical ultracentrifugation. **A&B. Sedimentation equilibrium analytical ultracentrifugation (SE-AUC).** Top panels show radial absorbance data (symbols) collected at three denoted rotor speeds fitted to single-species model (lines); lower panels show residuals from the model fit. With both particles, a singles species fit consistent with a DNA-wrapped octameric nucleosome was determined. Expected masses and calculated partial buoyant densities are provided in **Supplemental Table 3**, and masses derived from this analysis are shown in **Supplemental Table 5**. **C. Sedimentation velocity analytical ultracentrifugation (SV-AUC).** Van Holde-Weischet analysis of SV-AUC data from H3-601 (green), H3-αSat (cyan), and CENP-A-αSat (light red) NCPs at 25°C. Results were normalized for s_20,w_. Vertical profiles distinguish homogenous species from sloping distributions which indicated heterogeneity. The black arrow denotes bound fractions from CENP-A-αSat that indicate heterogeneity. **D. Dynamic Light Scattering (DLS) of α-satellite NCPs.** Panel shows the distribution of particles of H3-αSat (light red) and CENP-A-αSat (cyan) as a function of hydrodynamic radius, normalized for volume. The measured radius for the H3-αSat particle was 43.2 Å vs an effective radius of 50.8 Å for CENP-A-αSat.

Having confirmed the compositional homogeneity of particles on a biophysical level, we next interrogated the solution properties of these particles using sedimentation velocity analytical ultracentrifugation (SV-AUC). The experiment is performed at higher rotor speeds (and hence g forces) and in conditions known to be optimal for nucleosome integrity (*e.g.* low ionic strength, ambient temperature). The sedimentation coefficient is a hydrodynamic parameter that is sensitive to the size, shape, and density of particles in solution (Svedberg and Pedersen 1940). A model-independent van Holde-Weischet analysis (Vanholde and Weischet 1978) was employed to evaluate the monodispersity of these particles with minimal diffusion effects. The sedimentation properties observed by SV-AUC for the H3-601 particle agreed well with its atomic structure (s_20,w_ of 11.7 ± 0.9 for H3-601, representative of three independent trials, versus a calculated value of 10.9)(**Figure 2C**). Similarly, H3 histone octamer wrapped in α-satellite DNA displayed vertical profile by this analysis indicative of monodispersity (s_20,w_ of 11.2 ± 0.2, representative of three independent trials). However, while the average s value obtained was similar (s_20,w_ of 10.9 ± 0.2), the CENP-A-αSat particles displayed more polydispersity as evidenced by broader s distributions relative to the other particles assessed. Taken together with our SAS analyses, these data further indicate that the differences in solution properties observed by SAXS and AUC map not to differences in stoichiometry but physical differences in shape.

### The reversible assembly of NCPs under high pressure

While polydispersity for CENP-A-αsat NCPs in these SV-AUC experiments relative to other NCPs is apparent, that property was less pronounced in other complementary measures. A factor that distinguishes SV-AUC from these other methods employed is the occurrence of hydrostatic pressure under conditions of high centrifugal force. At the bottom of a SV-AUC cell at 40,000 RPM, calculated pressures upwards of ∼1.7 MPa are predicted (Schuck 2016) (whereas standard atmospheric pressure is ∼0.1 MPa). It has already been shown that histone octamers in isolation are sensitive to hydrostatic pressure (Silva, Villas-boas, and Clegg 1993; Scarlata, Ropp, and Royer 1989). To more directly investigate the possibility that pressure can affect NCP structure in solution, we turned to the high-pressure SAXS (HP-SAXS) resource at the Cornell High Energy Synchrotron Source (XBio Beamline) (Gillilan 2022). Hydrostatic pressure, when systematically applied, is a robust and powerful tool to explore reversible changes in macromolecular structure without the need for chemical excipients or mutations (Silva et al. 2014). Here, the goal was to better understand the differences in NCP structure and stability on authentic α-satellite DNA for H3 and CENP-A nucleosomes as a function of applied pressure.

We first examined the pressure-induced changes in H3-αSat structure at room temperature in 100 MPa increments, starting at 5 MPa and arriving at 300 MPa, followed by cycling between the two extremes (**Figure 3A**). At each pressure, the samples were allowed to equilibrate for five minutes before SAXS data collection. The structural properties obtained from this analysis are summarized in **Table 1** and the primary data shown in **Figure 3B**. As pressure was incrementally applied, changes in R_g_ and D_max_ were readily observed. Concomitant with the increases in R_g_ and D_max_, dimensionless Kratky plot analysis shows a steady transition away from a compact macromolecule to a more distended polymer as pressure increased, suggestive of unwrapping and disassembly of the NCP (**Figure 3C**, **Table 1**). To visualize these changes in shape, we performed *ab initio* electron density calculations using the program DENSS at each condition of pressure for both particles (**Figure 3D**). In the H3-αSat particles, the oblate ellipsoidal character of the initial particles at ambient pressure are modestly retained up to 200MPa. The simplest interpretation of these data is that the progressive unwrapping of the DNA ends of the NCPs increase the spatial extent and ellipsoidal character of the particles. At 300 MPa the particles undergo a dramatic shift to larger R_g_ and D_max_ values, indicating a more dramatic unfolding (**Table 1**). At 300 MPa both particles undergo a dramatic shift to a larger R_g_ and D_max_ (**Figure 4**). Pressure cycling directly between 300 MPa and 5 MPa was performed at the same time intervals and compared to the initial scattering profiles. Strikingly, the particle reverted to nearly its initial state, although the recovered R_g_ and D_max_ were always slighter higher for both particles, suggestive of hysteresis behavior: the “swelling” observed can be attributed to pressure-induced hydration effects, where interatomic contacts were replaced with water molecules, causing incorrect reassembly and folding.(Silva et al. 2014) This is further supported by the disparities observed in P(r) analysis from the same data, including (D_max_).

**Figure 3.**
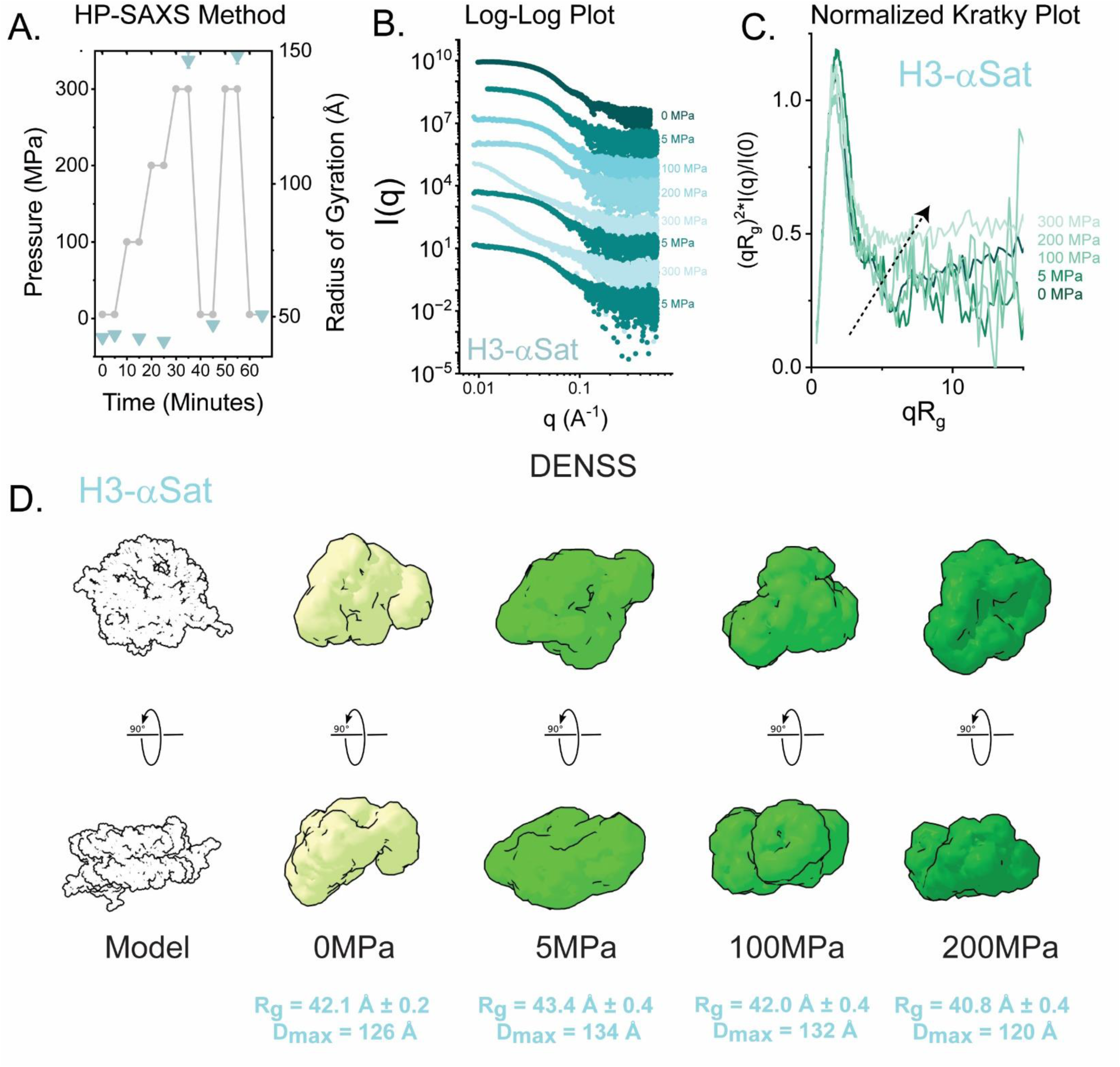
High-Pressure SAXS reveals reversible deformation of H3 NCPs assembled with α-satellite DNA. A. High pressure strategy (grey line and closed circle) is shown, with recorded R_g_ as determined by GNOM analysis at each pressure point shown as a cyan closed triangle. B. Pressure-dependent SAXS profiles from H3-αSAT, scaled arbitrarily along the y-axis. C. Normalized Kratky Plot analysis, showing the changes in the particle structure as a function of pressure. At the highest pressure, the particle retains some degree of folded character. D.DENSS analysis of α-satellite NCPs as a function of pressure. Shown in orthogonal views are DENSS reconstructions of H3-αSAT at different pressures, showing an oblate ellipsoidal character preserved through to the highest pressure. For reference, a canonical H3-601 atomic model is shown. R_g_ and D_max_ as determined by GNOM analysis are provided for each pressure point (**Table 1**).

**Figure 4.**
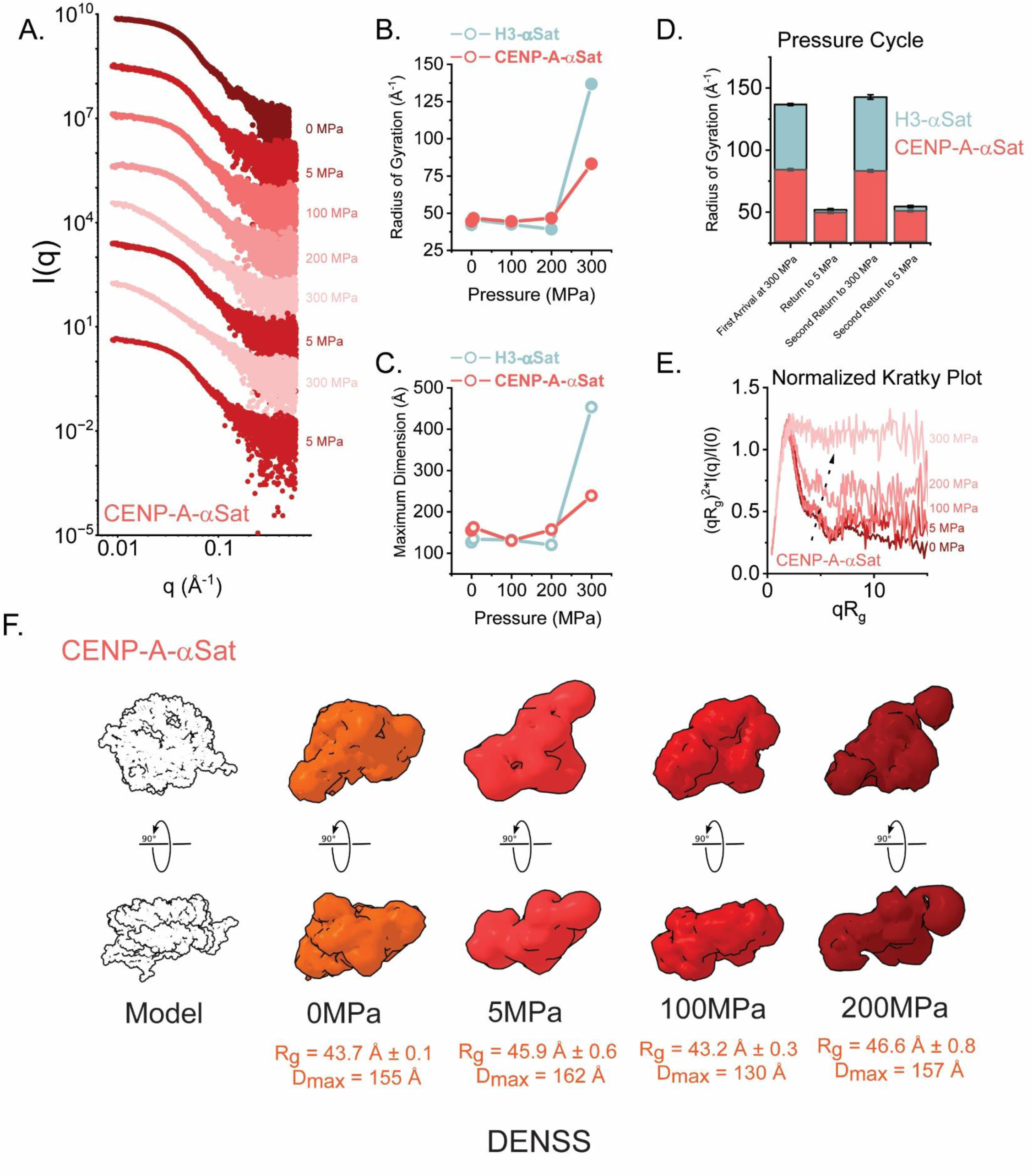
High-Pressure SAXS of CENP-A NCPs assembled with α-satellite DNA reveals distinct behaviors upon reversible deformation relative to H3 NCPs. **A.** Pressure-dependent SAXS profiles from CENP-A-αSat, scaled arbitrarily along the y-axis. **B&C.** Variation in R_g_ and maximum dimension as a function of pressure for H3-αSat (light red) and CENP-A-αSat (cyan). **D**. Stacked bar graph showing the reversible behavior under pressure for two iterations of changes in pressure from 300 MPa to 5 MPa. H3-αSat particles (shown in cyan) show a greater change in spatial extent versus the CENP-A-αSat particle (light red). **E.** Normalized Kratky Plot analysis, showing the changes in the particle structure as a function of pressure. At the highest pressure and unlike the H3-αSAT particle, CENP-A-αSat particle loses all folded character. **F.** DENSS analysis of CENP-A-α-Sat NCPs as a function of pressure. Shown in orthogonal views are DENSS reconstructions of CENP-A-αSAT at different pressures. For reference, a canonical H3-601 atomic model is shown. A transition from an oblate to elongated prolate particle is observed. R_g_ and D_max_ as determined by GNOM analysis are provided for each pressure point (**Table 1**).

### CENP-A nucleosomes also reversibly disassemble under pressure but display different properties at the most extreme pressures

Using the same pressure method, we next examined the CENP-A-αSat particle (**Figure 4A**). In contrast to the H3 result, we observe somewhat larger increases in R_g_ and D_max_ when pressure was applied through 200 MPa (**Figure 4B&C**), consistent with the unwrapping of the DNA. However, at 300 MPa the CENP-A particle adopts a much more compact structure than H3 nucleosomes (**Figure 4D**). Yet surprisingly, normalized Kratky plot analysis indicates the transition to an entirely unfolded polymer at 300 MPa. This contrasts with the H3-αSat result where some evidence of globularity is retained (Figure 3C). In agreement with these observations, DENSS analysis of the CENP-A-αSat particle data displays a more dramatic transition to a prolate ellipsoid form as pressure is increased (**Figure 4F**). For both particles, the observed I_0_ at the final pressure cycle back to the 5 MPa measurement condition suggests that sample mass was preserved throughout the experiment.

## Discussion

Our experiments revealed several similarities and differences between the solution behavior of conventional nucleosomes and their centromeric counterparts. SAXS and SANS performed at ambient pressure provide a view, albeit at lower resolution than crystallography or cryo-EM, of a generally shared NCP architecture in solution. Differences, potentially attributable to pressures under centrifugal force that exceed ambient pressure by more than an order of magnitude, are measurable via AUC between nucleosome types. NCP distortions under very high pressure (two additional orders of magnitude higher than in the AUC) lead to the most pronounced differences between nucleosomes containing the canonical histones versus histones where CENP-A replaces conventional histone H3.

The high degree of reversibility of NCP distortion at high pressures is very surprising and will stimulate further use HP-SAXS for studies of a variety of types of nucleosomes (and perhaps other chromatin complexes). The increase in R_g_ and D_max_ with increasing pressure correlates with a type of deformation and denaturation of the particle that is to be expected, but the ability of the particles to return to almost their original 3D shape indicates that the NCPs are robust, energetically optimized protein/DNA structures with a stable and self-directed structure. Of note, the magnitude of R_g_s observed at 300 MPa is potentially reminiscent of the magnitude observed by salt-induced dissociation of nucleosomes previously reported by time-resolved SAXS (TR-SAXS) measurements on NCPs, where protein signal was contrasted away with 50% sucrose and disassembly is monitored (Chen et al. 2017; Chen et al. 2014). The R_g_ and D_max_ observed for the H3-αSat particle at this extreme is entirely consistent with what would be calculated for a fully extended 147 bp B-form DNA using Flory’s Law (Caillet and Claverie 1974) (R_g_ of ∼144 Å, D_max_ of ∼500 Å). While our experiments do not leverage sucrose for total protein matching, we still expect the signal to still be dominated by the DNA scattering (Δρ_DNA_ of 220 e^−^/nm^3^ vs Δρ_protein_ of 90 e^−^/nm^3^). The difference in high pressure-induced deformation at 300 MPa between H3 NCPs and CENP-A NCPs has two seemingly contradictory findings: CENP-A has a more extended shape but smaller R_g_ and D_max_. A potential explanation is that CENP-A NCPs completely but reversibly dissociate some histone subunits at very high pressures, while H3 NCPs remain octameric at high pressures.

High pressure studies are directly relevant to considerations of deep-sea life. Deep-sea life comprises a major percentage of the planet’s total biomass (Bar-On, Phillips, and Milo 2018), and includes species of prokaryotes and archaea with chromatin-like assemblies (Yamaguchi et al. 2012; Henneman et al. 2018; Takai and Horikoshi 1999) and eukaryotes that have adapted their chromatin to function at extreme temperatures (both near 0°C and exceeding 50°C), pressures, and ionic strength (dissolved salts upwards of ∼1 M)(Gage and Tyler 1991). Little is known about this aspect of deep-sea life on a biochemical, biophysical, and structural level, and such insights would inform our broader understanding of chromatin structure and dynamics across all species of life. HP-SAXS is uniquely poised to directly interrogate the physical properties of these biological assemblies in native-like conditions. The pressures achievable at the Cornell CHESS SAXS resource well exceed those pressures encountered by deep sea life at up to ∼10 km depths (∼100 MPa), providing the opportunity to probe the stability, disassembly, and reassembly of chromatin and chromatin-like structures as was performed here in this study, and to determine how nature has evolved these assemblies to persist in extreme conditions.

In closing, we highlight that nucleosomes in a chromosome undergoing biological processes are encountering forces that may (or are known to) distort and dissociate them. These include DNA metabolic processes (replication and transcription), chromatin disassembly and remodeling by ATPases, spindle forces at cell division, nuclear deformation in tissues under mechanical stress, and more. The methodologies and findings described in this study provide further approaches and understanding into this challenging aspect of chromosome studies.

## Methods

### Preparation of reconstituted mononucleosomes

Reconstituted mononucleosomes were prepared as previously described using recombinant histones and DNAs (Sekulic and Black 2016). The following DNA sequences were used:

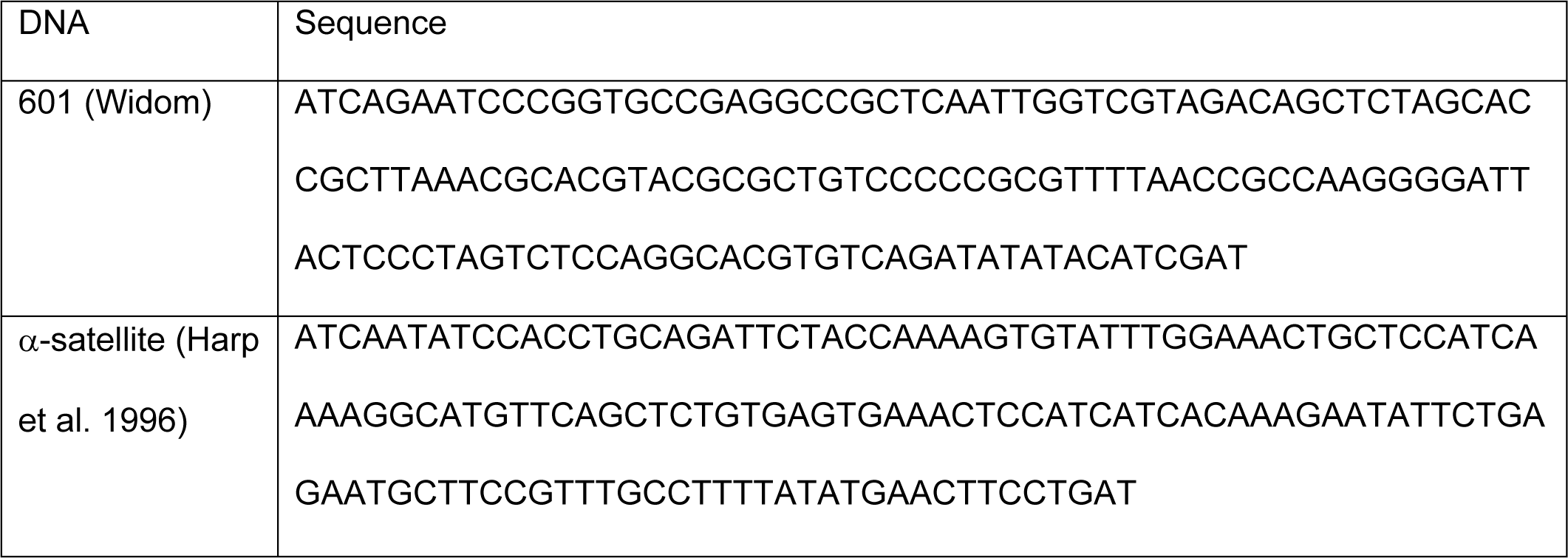

Reconstituted nucleosomes were uniformly positioned on sequences by thermal shifting (heating at 55°C for two hours). After this, nucleosomes were separated from free DNA, nucleosomes with alternative positions on the DNA and higher-order aggregates using 5% preparative native gel electrophoresis (PrepCell 491, (37 mm diameter x 7 cm height), Bio-Rad, Hercules, California, USA). The histone composition and stoichiometry were verified with 2D-PAGE. Finally, samples were dialyzed into 20 mM potassium cacodylate pH 7.0 and 1 mM EDTA for all SANS experiments or 20 mM Tris-HCl pH 7.5, 1 mM EDTA, and 1 mM DTT for all other biophysical analyses described herein. Sample concentrations were determined using Bradford Assay (Bradford 1976) and by measuring absorbance at 260nm (DNA) in clean nucleosome samples.

### Dynamic Light Scattering

Samples at 0.3 mg/mL concentration were analyzed using a Nanobrook Omni particle sizer (Brookhaven Instruments Corporation, Holtsville, NY, USA). Data were recorded at 25°C in polystyrene 1-cm cells using a standard diode laser at 640 nm, with scattering recorded at an angle of 90°. Three scans were recorded for each sample and hydrodynamic radii (Stokes radii, R_s_) were calculated using the BIC Particle Solutions software v3.6.0.7122.

### Sedimentation Equilibrium (SE) and Sedimentation Velocity (SV) Analytical Ultracentrifugation

Analytical ultracentrifugation experiments were performed with an XL-A analytical ultracentrifuge or Optima (Beckman-Coulter, Indianapolis, IN, USA) and a TiAn60 rotor with six-channel (for SE) or two-channel (for SV) charcoal-filled epon centerpieces and quartz or sapphire windows. SE data were collected at 4°C with detection at 260 nm for three sample concentrations. SE analyses were carried out using global fits to data acquired at multiple speeds for each concentration with strict mass conservation using the program SEDPHAT (25). Error estimates for masses derived with mass-averaged partial specific volumes (ʋ_bar_) were determined from a 1,000-iteration Monte Carlo simulation. A partial specific volume value for the different particles examined were calculated by the program MuLCH (Whitten, Cai, and Trewhella 2008a) based on chemical composition.

Complete SV profiles were recorded on samples (OD_260nm_ of 0.5-1.0) every 30 seconds at 260 nm for 50-200 boundaries at 26,000 rpm. Selected boundaries from the dataset were analyzed using the program SEDFIT to generate van Holde-Weischet plots. Solvent density was determined gravimetrically at room temperature (d = 1.01 g/mL ± 0.01 g/mL), and a viscosity of η = 0.001 poise was used in all analyses.

### Experimental Considerations for SAXS Analysis

Several experimental considerations were made and optimized for this study. The intensity of scatter from a particle can be expressed as:

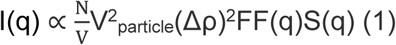

where 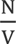 is the number of proteins per unit volume (concentration), V_particle_ is the volume of the individual particle, Δρ the contrast, *FF(q)* is the form factor, or scattering component from a single particle rotationally averaged, and *S(q)* is the interparticle structure factor.

Because of the relatively electron-rich nucleic acid component comprising the bulk of this particle’s mass and exterior and the particles relatively large size, we found that relatively low sample concentrations of particle (∼0.75 mg/mL-1.5 mg/mL) provided measurable scatter at both synchrotron sources and with a rotating anode X-ray source, and with experimental neutron sources, as intensity of scattering varies as the square of volume (eqn. 1)). The added benefit of working with samples at these relatively dilute concentrations is the minimization of any potential interparticle interference (*S(q)*) that could confound structural analysis.

### Small-angle X-ray scattering (SAXS) Data Collection

X-ray scattering data were measured on at two different synchrotron sources: beamline CHEX at the Cornell University High Energy Synchrotron Source (Acerbo, Cook, and Gillilan 2015; Skou, Gillilan, and Ando 2014) (CHESS Ithaca, NY) and beamline SIBYLS at the Advanced Light Source (Hura et al. 2009). Data were also recorded using a rotating anode SAXS instrument as described previously. In all cases, the forward scattering from the samples studied was recorded on a CCD or multiwire detector and circularly averaged to yield one-dimensional intensity profiles as a function of *q* (*q* = 4πsinθ/λ, where 2θ is the scattering angle). Samples were centrifuged at 10,000 × g for three min at 4°C prior to 0.5 s – 20 s X-ray exposures at 20°C. Scattering from a matching buffer solution was subtracted from the data and corrected for the incident intensity of X-rays. Replicate exposures were examined carefully for evidence of radiation damage by Guinier analysis and Kratky plot analysis. Silver behenate powder was used to locate the beam center and to calibrate the sample-to-detector distance. All the preparations analyzed were monodisperse, as evidenced by linearity in the Guinier region of the scattering data (where *qR_g_* ≤ 1.2) and agreement of the I_0_ and R_g_ values determined with inverse Fourier transform analyses using the program GNOM (Svergun 1992). Experimental details unique to each X-ray source are provided in Supplemental Methods.

### Small-angle neutron scattering (SANS) Data Collection

Neutron scattering data were measured at two different research reactor locations: beamline NG-3 of the National Institutes of Standards and Technology (NIST) Center for Neutron Research (Glinka et al. 1998), and beamline CG-3 of the Oak Ridge National Laboratories High Flux Isotope Reactor (HFIR) (Heller et al. 2008). Experimental details unique to each beamline are provided in Supplemental Methods. Samples were prepared by dialysis at 4°C against matching buffers (20 mM potassium cacodylate pH 7.0 and 1 mM EDTA) containing 0%, 20%, 70%, 80%, or 95% D_2_O for a minimum of three hours across a 6-8 kD cutoff membrane (D-tube dialyzer (Novagen)). Samples were centrifuged at 10,000 x g for three min at 4°C and then loaded into Hellma quartz cylindrical cells (outer diameter of 22 mm) with either 2-mm (for 95% and 80% D_2_O) or 1-mm pathlengths (70%, 20%, and 0% D_2_O). Before and during the experiment the samples were maintained at 6°C. Sample concentrations for the SANS measurements were determined by Bradford analysis and are shown in Table 2. At both locations, scattering neutrons were detected with a two-dimensional position-sensitive detector and data reductions are performed using beamline-specific software. Raw counts were normalized for incident intensity and corrected for empty cell counts, ambient room background counts, and non-uniform detector response. Data were placed on an absolute scale and radially averaged to produce one-dimensional scattered intensity *I(q)* versus *q* profiles. Data collection times varied from 0.5 to 5 hours depending on the instrument configuration, sample concentration, and amount of D_2_O present in the sample. Multiple sample-to-detector distances were employed, and data were merged to create the final scattering profile for data analysis. At both locations, a wavelength of 6 Å and with a spread of 0.15 was employed. We observed good agreement between R_g_ and I_0_ determined from inverse Fourier analysis using GNOM and that determined by classical Guinier analysis (Guinier 1939). The program MuLCH (Whitten, Cai, and Trewhella 2008b) was used to calculate theoretical contrast and to initially analyze contrast variation data, assuming ∼50% proton exchange based on previously reported hydrogen-deuterium mass spectrometry studies (Sekulic et al. 2010). Stuhrmann plot analyses were performed manually using Origin version 2024b (Originlab Corp., Northampton, MA, USA). All of the preparations analyzed were monodisperse, as evidenced by linearity of sqrt(I_0_/c) versus fractional D_2_O plots (See Supplemental Figure 2) and by comparison of the linear Guinier region of the scattering data with the I_0_ and R_g_ values determined with inverse Fourier transform analysis by the programs GNOM (Semenyuk and Svergun 1991) (Table 2). Additional experimental details specific to each location are provided in Supplemental Methods.

### Molecular Mass Calculations from Contrast Variation SANS Data

The scattered intensities from the protein-DNA complexes were decomposed into the scattering from their components, I_PROT_(q) and I_RNA_(q), using the equation (Kuzmanovic et al. 2003):

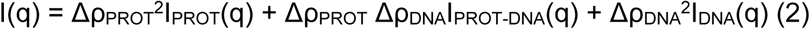

where Δρ = (ρ − ρ_s_) is the contrast, or the difference between the scattering length density of the molecule (ρ) and the solvent (ρ_s_). The cross-term, I_PROTDNA_(q), represents the interference function between the protein and DNA components. The known quantities in equation 1 are Δρ_PROT_ and Δρ_RNA_ and the unknowns are *I_PROT_(q), I_DNA_(q)*, and *I_PROTDNA_(q)*. Since measurements were made at four-five different contrasts, or D_2_O/H_2_O buffer conditions, there is sufficient information to solve for the three unknown component intensities from the set of simultaneous equations for *I(q)* at each contrast:

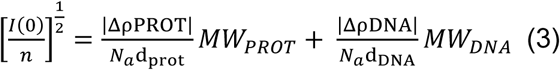

Where N_a_ is Avogadro’s number, Δρ is the calculated net scattering length density, and d is the mass density, where d_prot_ = 1.35 g/cm^3^ and d_DNA_ = 1.69 g/cm^3^. The I_0_ values in absolute units (cm^−1^) obtained from the GNOM analysis of the data for each D_2_O/H_2_O buffer are used with the measured number densities to solve the set of simultaneous equations for these two unknowns to obtain the MW values for the protein and DNA components (MW_PROT_ and MW_DNA_) separately in the nucleosome particle, where the total MW value is then the sum of the two component MW values. Our contrast calculations assume 50% of acidic proteins accessible by solvent (Sekulic et al. 2010).

### Molecular Modeling

Complete atomistic models of the canonical nucleosome are derived from the crystal structure of canonical nucleosome on the 145 bp long Widom 601 sequence (PDB ID 3LZ0 (Kato et al. 2011)). The model the CENP-A nucleosome was derived from the crystal structure of CENP-A nucleosome on the 147 bp engineered palindromic α-satellite DNA (PDB ID 3AN2). For modeling, the DNA sequence was mutated to the 145 bp Widom 601 sequence. Missing sequences at the N and C-termini of the respective histone components were modelled as unstructured coils with known amino acid sequences. The NAMD (Phillips et al. 2020) program employing CHARMM43 forcefields was used to perform molecular dynamics. The resulting model was gradually relaxed by energy minimization and subsequent simulation in a box of water. Tails alone were relaxed further *in vacuo* for 1000 fs to collapse their position. The models shown were rendered using the program PyMOL 2.5.2 Molecular Graphics System (Schrodinger, LLC, New Your, NY). Hullrad (Fleming and Fleming 2018) and WinHYDRPRO (Garcia De La Torre, Huertas, and Carrasco 2000) were used to calculate the predicted hydrodynamic properties of these atomic models.

### High Pressure Small-Angle X-ray Scattering (HP-SAXS)

HP-SAXS experiments were conducted on beamline ID7A1 at the Cornell High Energy Synchrotron Source (CHESS) using a hydrostatic pressure cell with a maximum operating pressure of 400 MPa (Acerbo, Cook, and Gillilan 2015; Skou, Gillilan, and Ando 2014). Samples were prepared at a concentration of 1.2 mg/mL and centrifuged at 10,000 x g for three minutes at 4°C prior to measurement. The sample (40 μL) was filled into a disposable acrylic polymethyl methacrylate (PMMA) sample cell and sealed with high-vacuum grease (Dow Corning, Midland, MI, USA), which acted as a freely moving piston to equilibrate the sample to the surrounding pressure medium (water). The HP-SAXS cell design has been described in the literature (Rai et al. 2021).

Hydrostatic pressure cycles were performed at 25°C and between ambient pressure (0 MPa) to 300 MPa in increments of 100 MPa. At each pressure, the sample was allowed to equilibrate for five minutes before collecting the SAXS data. The beam was blocked during equilibration to prevent radiation damage. For individual measurements, the sample was exposed for a total of 10 s (10 exposures of 1s each). The matched buffer blanks were measured at identical pressures for proper background subtraction. The photon energy of the X-ray beam was 14.09 keV (0.8788 Å) at 1.6 × 10^11^ photons/second with a standard beam size (250 μm × 250 μm). The data were collected using an EIGER X 4 M detector (DECTRIS, Switzerland) with a pixel size of 75 μm × 75 μm and an active area of 155.2 × 162.5 mm. The sample-to-detector distance was 1.772 m, with the SAXS detector covering a collected *q* range of 0.0083 Å^−1^ < *q* < 0.6925 Å^−1^. The wavevector is defined as *q* = (4π/λ) sin θ, where 2θ is the scattering angle and λ is the wavelength of the incident radiation (**Table 1**). All data were reduced and analyzed using the program RAW (Hopkins 2024).

### *Ab initio* electron density reconstruction using DENSS

DENSS (Grant 2018a) was used to calculate the *ab initio* electron density directly from GNOM (Svergun 1992) outputs, as implemented in the program RAW. Twenty reconstructions of electron density were performed in the slow mode with default parameters and subsequently averaged and refined with no symmetry restraints. Reconstructions were visualized using either UCSF ChimeraX (Meng et al. 2023), or PyMOL 2.5.2 Molecular Graphics System (DeLano) (Schrodinger, LLC, New Your, NY) with five contour levels of density rendered with these respective colors: 15σ (red), 10σ (green), 5σ (cyan), 2.5σ (blue), and −0.7σ (blue).The sigma (σ) level denotes the standard deviation above the average electron density value of the generated volume.

## Acknowledgements

We thank Steven Stayrook (Penn), and Qian Shuo and Sai Pengali (ORNL) for their technical assistance. We thank Jochen Hub (Georg-August-University, Germany) for adjustments made to the WAXIS server to accommodate our structural models. We also thank Thomas Grant (University of Buffalo) for helpful discussions regarding the application of DENSS.

## Author Contributions

Conceptualization: Kushol Gupta, Nikolina Sekulić, Gregory D. Van Duyne, Ben E. Black; Methodology: Kushol Gupta, Nikolina Sekulić, Susan Krueger, Joseph Curtis, Richard E. Gillilan, Reidar Lund; Formal analysis and investigation: Kushol Gupta, Nikolina Sekulić, Praveen Kumar Allu, Nicklas Sapp, Qingqiu Huang, Kathryn Sarachan, Mikkel Christensen, Susan Krueger^;^ Writing - original draft preparation: Kushol Gupta, Nikolina Sekulic, Ben E. Black; Writing - review and editing: Kushol Gupta, Nikolina Sekulić, Praveen Kumar Allu, Nicklas Sapp, Qingqiu Huang, Kathryn Sarachan, Susan Krueger, Joseph Curtis, Richard E. Gillilan, Reidar Lund, Gregory D. Van Duyne, Ben E. Black; Funding acquisition: Gregory D. Van Duyne, Ben E. Black. All authors read and approved the final manuscript.

## Data Availability

Several structure coordinates available in the PDB database were used in the present studies, which can be located under accession numbers: PDB ID 3LZ0 (Kato et al. 2011) and PDB ID 3AN2 (Tachiwana et al. 2011), X-ray and Neutron scattering profiles will be made available at the SASDB upon publication. All other data will be made available upon request.

## Ethical approval

Not Applicable.

## Consent to participate

Not Applicable.

## Consent for publication

Not Applicable.

## Supplemental Methods

### Rotating Anode

Preliminary data were recorded on a Rigaku PSAXS small-angle X-ray scattering system equipped with Osmic mirror optics, a three-pinhole enclosed pre-flight path, an evacuated sample chamber with customized sample holder maintained at 4°C, and a gas-filled multi-wire detector. The instrument is served by a Rigaku MicroMax-007 HF microfocus rotating anode generator (Rigaku America, Woodland, T.X., U.S.A.). The forward scattering from the samples studied was recorded on a CCD detector and circularly averaged to yield one-dimensional intensity profiles as a function of *q* (*q*=4πsinθ/λ, where 2θ is the scattering angle, in units of Å^−1^). Data were reduced using SAXSGui v2.05.02 (Rigaku America) and matching buffers were subtracted to yield the final scattering profile. The sample-to-detector distance and beam center were calibrated using silver behenate and intensity converted to absolute units (cm^−1^) using a known polymer standard.

### Small-Angle X-ray Scattering at the Advanced Light Source Beamline 12.3.1 (SIBYLS)

Samples were centrifuged at 3,000 rpm for 10 min at 4°C prior to data collection. Data was collected using a 96-well plate handling sample robot, as previously described (Hura et al. 2009). All samples were characterized with 0.5, 1, and 6 s exposures at 20°C, at a wavelength of 1.0 Å. Data were automatically reduced using custom software to provide one-dimensional intensity profiles as a function of *q* (*q*=4πsinθ/λ, where 2θ is the scattering angle). Accessible scattering was recorded in the range of 0.010 < *q* < 0.35 Å^−1^.

### Small-angle Neutron Scattering at National Institutes of Standards and Technology Center for Neutron Research NG-3 (Glinka et al. 1998)

Samples were prepared by dialysis at 4°C against matching buffers containing 20 mM potassium cacodylate pH 7.0, 5 mM EDTA and 0%, 20%, 70%, 80%, or 95% D_2_O for a minimum of three hours across membrane with 6-8kD cutoff (D-tube dialyzer (Novagen)). Samples were centrifuged at 10,000 *g* for 3 min at 4°C and then loaded into Hellma quartz cylindrical cells (outside diameter of 22 mm) with either 2-mm (for 95% and 80% D_2_O) and 1-mm pathlengths (70%, 20%, and 0% D_2_O) and maintained at 6°C. Sample concentrations for the SANS measurements were determined by Bradford analysis(Bradford 1976) and are shown in Table 2.

Scattered neutrons were detected with a 64 cm × 64 cm two-dimensional position-sensitive detector with 128 × 128 pixels at a resolution of 0.5 cm/pixel. Data reduction was performed using the NCNR Igor Pro macro package (Kline 2006). Raw counts were normalized to a common monitor count and corrected for empty cell counts, ambient room background counts and non-uniform detector response. Data were placed on an absolute scale by normalizing the scattered intensity to the incident beam flux. Finally, the data were radially-averaged to produce scattered intensity (*I(q)*, in cm^−1^) versus *q* (Å^−1^) profiles. The scattered intensities from the samples were further corrected for buffer scattering and incoherent scattering from hydrogen in the samples. Data collection times varied from 0.5 hour to 2 hours, depending on the instrument configuration, sample concentration and buffer conditions. Sample-to-detector distances of 11 m (*q*-range 0.006-0.043 Å^−1^, where *q*= 4πsin(θ)/λ, where λ is the neutron wavelength and 2θ is the scattering angle), 5 m (*q*-range 0.011–0.094 Å^−1^), and 1.5 m (detector offset by 20.00 cm, *q*-range 0.03–0.4 Å^−1^) were measured at a wavelength of 6 Å with wavelength spread of 0.15. We observed good agreement between R_g_ and I_0_ determined from inverse Fourier analysis using GNOM (Svergun 1992) and that determined by Guinier analysis. The program MuLCH (Whitten, Cai, and Trewhella 2008) was used to calculate theoretical contrast and to analyze contrast variation data.

### Small-angle Neutron Scattering (SANS) at Oak Ridge HFIR CG-3

Experiments were conducted at the CG-3 BioSANS instrument at Oak Ridge National Laboratory (ORNL, Oak Ridge, TN). The wavelength of 6.0 Å with a wavelength spread of 0.15 Å was utilized at a 6 m sample-to-detector distance for one-hour exposures, providing an accessible *q* (where *q*=4πsinθ/λ, where 2θ is the scattering angle, in units of Å^−1^) of 0.008 < *q* < 0.15. Data were recorded at 6°C for all measurements in 1-mm (70%, 20%, and 0% D_2_O) or 2-mm (for 95% and 80% D_2_O) Hellma quartz cylindrical cells. To obtain normalized scattering intensities *I(q)* (cm^−1^) as a function of *q* (Å^−1^), empty cell and buffer cell scattering were subtracted from the sample scatter and normalized to absolute intensity units using a known polymer standard. Data reduction was performed using customized reduction scripts with the Mantid platform (Arnold et al. 2014). The program MuLCH (Whitten, Cai, and Trewhella 2008) was used to calculate theoretical contrast and to analyze contrast variation data. All scattering data were analyzed by the inverse Fourier transform using the program GNOM assessed by classical Guinier analyses.

## Supplemental Tables

**Supplemental Table 1.** Calculated Masses of Particles studied using Linear analysis of SANS I_0_.

|  | Method 1 |  |  | Method 2 |  |
| --- | --- | --- | --- | --- | --- |
| Sample | M<sub>protein</sub> (kDa) | M<sub>DNA</sub> (kDa) | Total M<sub>Complex</sub> | M<sub>protein</sub> (kDa) | Total M<sub>Complex</sub> |
| CENP-A-601 | 131 ± 25 | 72 ± 23 | 203 ± 35 | 118 ± 19 | 207 ± 19 |
| H3-601 | 127 ± 12 | 86 ± 11 | 213 ± 16 | 125 ± 8 | 214 ± 8 |
| CENP-A-αSat | 77 ± 16 | 121 ± 20 | 198 ± 24 | 87 ± 16 | 176 ± 16 |
| H3-αsat | 76 ± 31 | 140 ± 39 | 216 ± 49 | 80 ± 11 | 169 ± 29 |

Method 1: M<sub>protein</sub> and M<sub>DNA</sub> were found by solving simultaneous equations using $I_0$ values from 4-5 SANS contrast points.
Concentrations were determined by Bradford assay.
Method 2: M<sub>DNA</sub> was fixed and M<sub>protein</sub> was determined using simultaneous equations.

**Supplemental Table 2.** Parameters derived from Stuhrmann Analysis of SAXS/SANS data.

| Sample | R<sub>c</sub> (Å) | $\alpha$ (cm<sup>-1</sup>) | $\beta$ (cm<sup>-2</sup>) | Fit R<sup>2</sup> |
| --- | --- | --- | --- | --- |
| **H3-601** | 39.0 ± 0.2 | 417.7 ± 81.2 | -698.6 ± 327.0 | 0.97 |
| *Hjelm 1977* | 40.5 | 450 |  |  |
| **CENP-A-601** | 39.9 ± 0.3 | 544.4 ± 153.5 | -320.1 ± 527.4 | 0.90 |
| **H3-αSat** | 40.9 ± 0.2 | 552.3 ± 159.4 | -596.0 ± 461.7 | 0.86 |
| **CENP-A-αSat** | 40.1 ± 0.4 | 305.9 ± 138.1 | -805.6 ± 452.6 | 0.97 |

**Supplemental Table 3.**
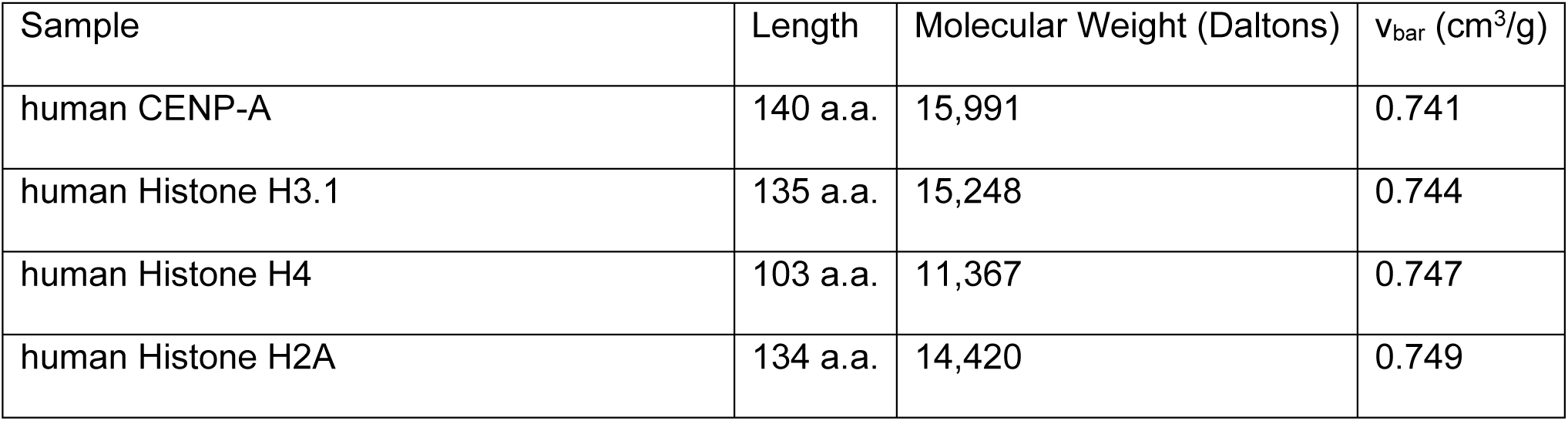
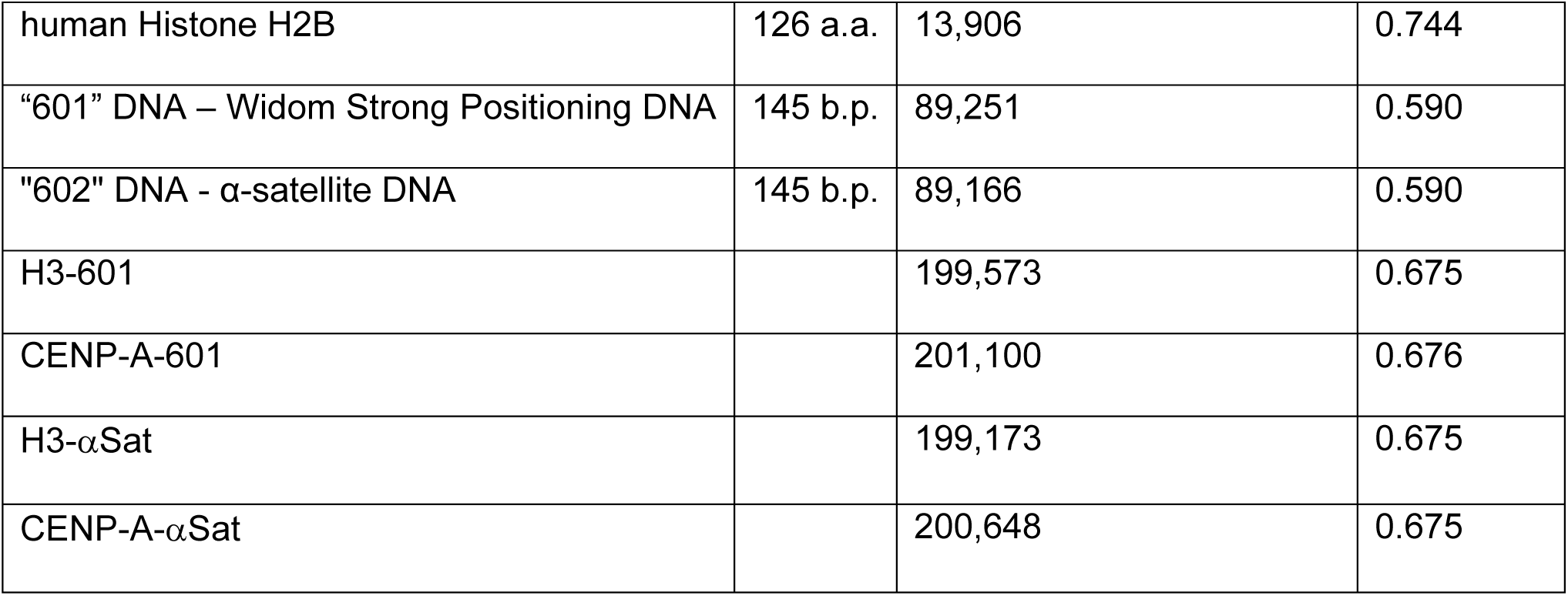
Calculated properties of proteins and DNAs used in this study.

| Sample | Length | Molecular Weight (Daltons) | v<sub>bar</sub> (cm<sup>3</sup>/g) |
| --- | --- | --- | --- |
| human CENP-A | 140 a.a. | 15,991 | 0.741 |
| human Histone H3.1 | 135 a.a. | 15,248 | 0.744 |
| human Histone H4 | 103 a.a. | 11,367 | 0.747 |
| human Histone H2A | 134 a.a. | 14,420 | 0.749 |
| human Histone H2B | 126 a.a. | 13,906 | 0.744 |
| "601" DNA – Widom Strong Positioning DNA | 145 b.p. | 89,251 | 0.590 |
| "602" DNA - α-satellite DNA | 145 b.p. | 89,166 | 0.590 |
| H3-601 |  | 199,573 | 0.675 |
| CENP-A-601 |  | 201,100 | 0.676 |
| H3-αSat | | 199,173 | 0.675 |
| CENP-A-αSat | | 200,648 | 0.675 |

**Supplemental Table 4.** Global Core-Shell Cylinder Fitting of SAXS/SANS Data using SASVIEW (related to Supplemental Figure 5).

| **Sample** | **Data** | **Fit Radius** | **Fit Length** | **$\chi^2$** |
| --- | --- | --- | --- | --- |
| H3-601 | Global Fit (5) | 19.5 Å ± 0.62 | 59.5 Å ± 2.8 | 2.8 |
|  | 0% D<sub>2</sub>O (SANS) |  |  | 1.6 |
|  | 20% D<sub>2</sub>O (SANS) |  |  | 2.4 |
|  | 80% D<sub>2</sub>O (SANS) |  |  | 0.9 |
|  | 90% D<sub>2</sub>O (SANS) |  |  | 1.1 |
|  | SAXS |  |  | 1.3 |
| CENP-A-601 | Global Fit (5) | 18.9 Å ± 0.12 | 70.3 Å ± 0.9 | 2.2 |
|  | 0% D<sub>2</sub>O (SANS) |  |  | 1.2 |
|  | 20% D<sub>2</sub>O (SANS) |  |  | 2.9 |
|  | 80% D<sub>2</sub>O (SANS) |  |  | 0.7 |
|  | 90% D<sub>2</sub>O (SANS) |  |  | 1.1 |
|  | SAXS |  |  | 0.6 |
| H3-αSat | Global Fit (6) | 18.9 Å ± 0.05 | 68.6 Å ± 0.3 | 3.7 |
|  | 0% D<sub>2</sub>O (SANS) |  |  | 1.1 |
|  | 20% D<sub>2</sub>O (SANS) |  |  | 0.9 |
|  | 70% D<sub>2</sub>O (SANS) |  |  | 3.4 |
|  | 80% D<sub>2</sub>O (SANS) |  |  | 2.5 |
|  | 90% D<sub>2</sub>O (SANS) |  |  | 2.8 |
|  | SAXS |  |  | 1.6 |
| CENP-A-αSat | Global Fit (6) | 18.1 Å ± 0.04 | 74.3 Å ± 0.2 | 2.6 |
|  | 0% D<sub>2</sub>O (SANS) |  |  | 1.1 |
|  | 20% D<sub>2</sub>O (SANS) |  |  | 0.8 |
|  | 70% D<sub>2</sub>O (SANS) |  |  | 1.5 |
|  | 80% D<sub>2</sub>O (SANS) |  |  | 0.9 |
|  | 90% D<sub>2</sub>O (SANS) |  |  | 0.9 |
|  | SAXS |  |  | 1.6 |

**Supplemental Table 5.** Parameters derived from Analytical Ultracentrifugation.

| Sample | Sedimentation Equilibrium (SE)(Daltons) |  |
| --- | --- | --- |
|  | 25°C | 4°C |
| H3-601 | 202,787 ± 1,437 | 187,233 ± 1,182 |
| H3-αSat | 199,226 ± 1,345 | 208,036 ± 1,316 |
| CENP-A-αSat | 197,476 ± 4,887 | 193,466 ± 4,339 |

SE-AUC experiments were carried out at 26,000 RPM in 20 mM Tris-HCl pH 7.5, 1 mM EDTA, and 1 mM DTT. Masses derived from SE experiments were calculated by global fitting of three concentrations of particle at three different speeds (7K, 9K, and 11Krpm).

## Supplemental Figures

**Supplemental Figure 1.**
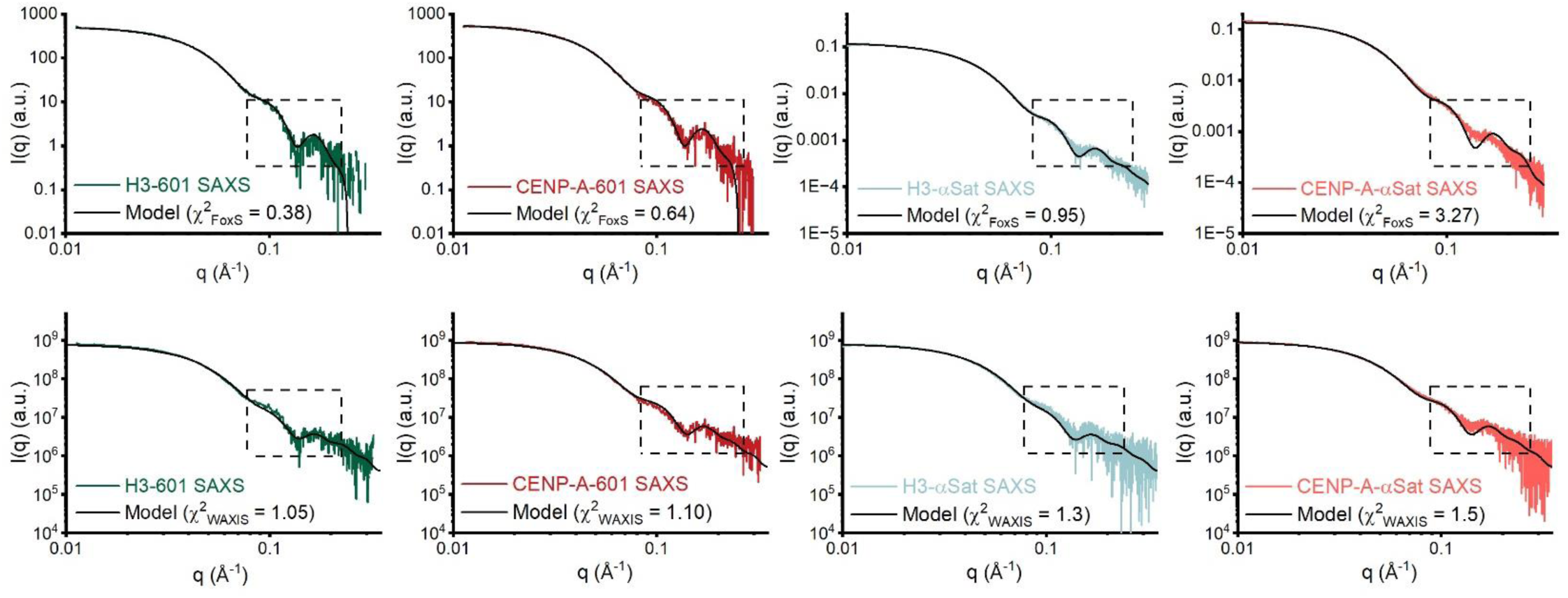
FoxS and WAXIS fitting of atomic models to SAXS Data. Shown are the respective model fits (black lines) to SAXS data performed using FoxS (upper panels, implicit solvent boundary model) or WAXiS (lower panels, explicit solvent boundary calculation) for H3-601 (green), CENP-A-601 (red), H3-αSat (cyan), and CENP-A-αSat (light red). Shown in black boxes are middle-q regions of the fit most discrepant in the fitting, with a χ^2^ for the fit provided in the respective graph legends.

**Supplemental Figure 2.**
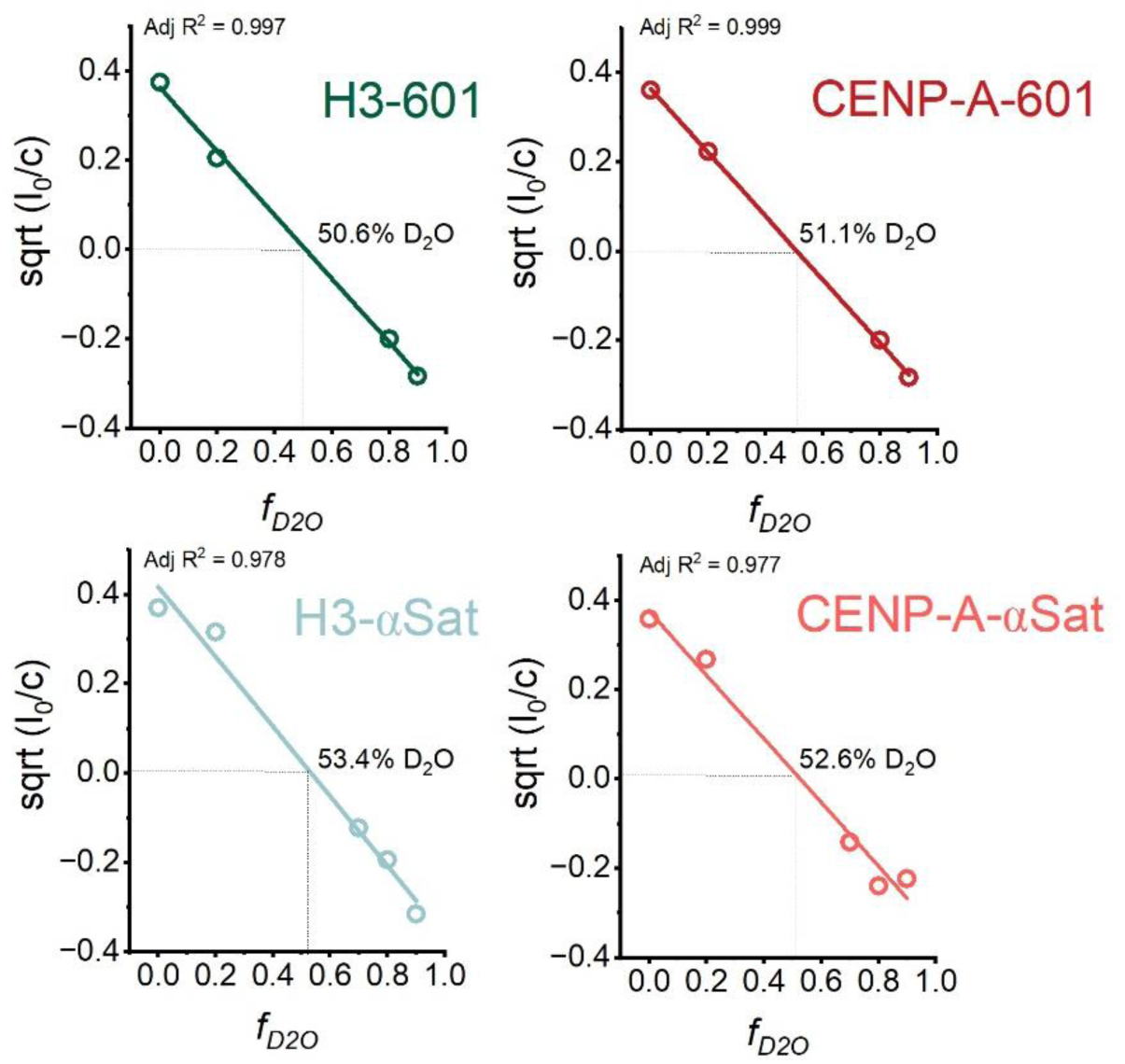
Zero-Angle Scattering from SANS. The contrast-dependence of the zero-angle scattering is shown. The square root of the extrapolated zero angle scatter (I_0_, in absolute units of cm^−1^) divided by concentration (c) in mg/ml, plotted against the fractional D_2_O (*f_D2O_*). The fractional D_2_O can be directly related to the solvent scattering length density for each contrast. A linear plot in all four cases indicates monodispersity in these preparations (Stuhrmann and Duee 1975) and the least squares fit provides the overall contrast point where intensity is zero.

**Supplemental Figure 3.**
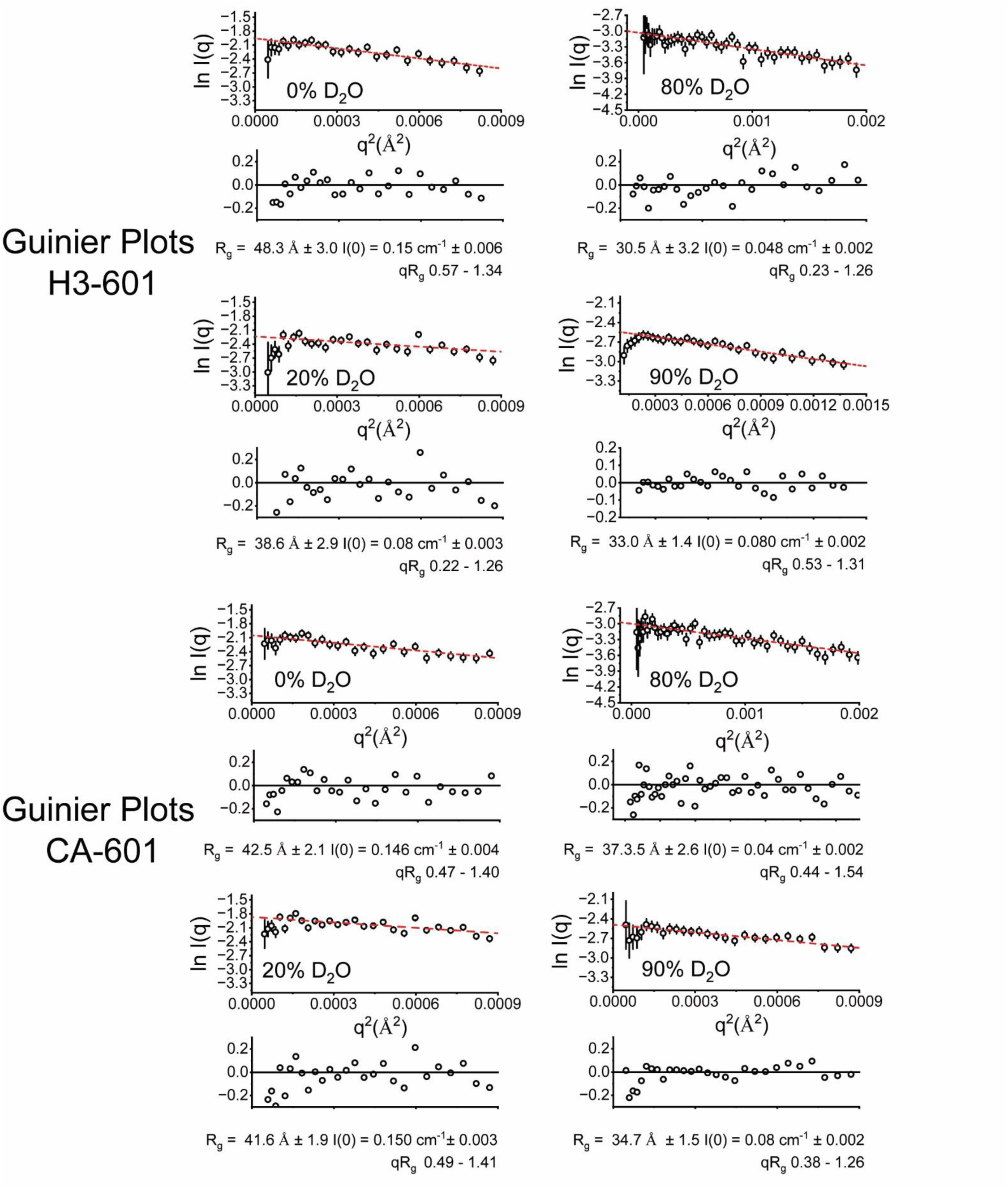
SANS Guinier Plots for H3-601 and CENP-A-601. Guinier plot analyses (ln (I) vs. q^2^) of SAXS data (black open circles) for H3-601 and CENP-A601 NCPs, with residuals from the fitted lines shown below. Monodispersity is evidenced by linearity in the Guinier region of the scattering data and agreement of the I_0_ and R_g_ values determined with inverse Fourier transform analysis by the programs GNOM (Table 2). Guinier analyses were performed where qR_g_ ≤ 1.4.

**Supplemental Figure 4.**
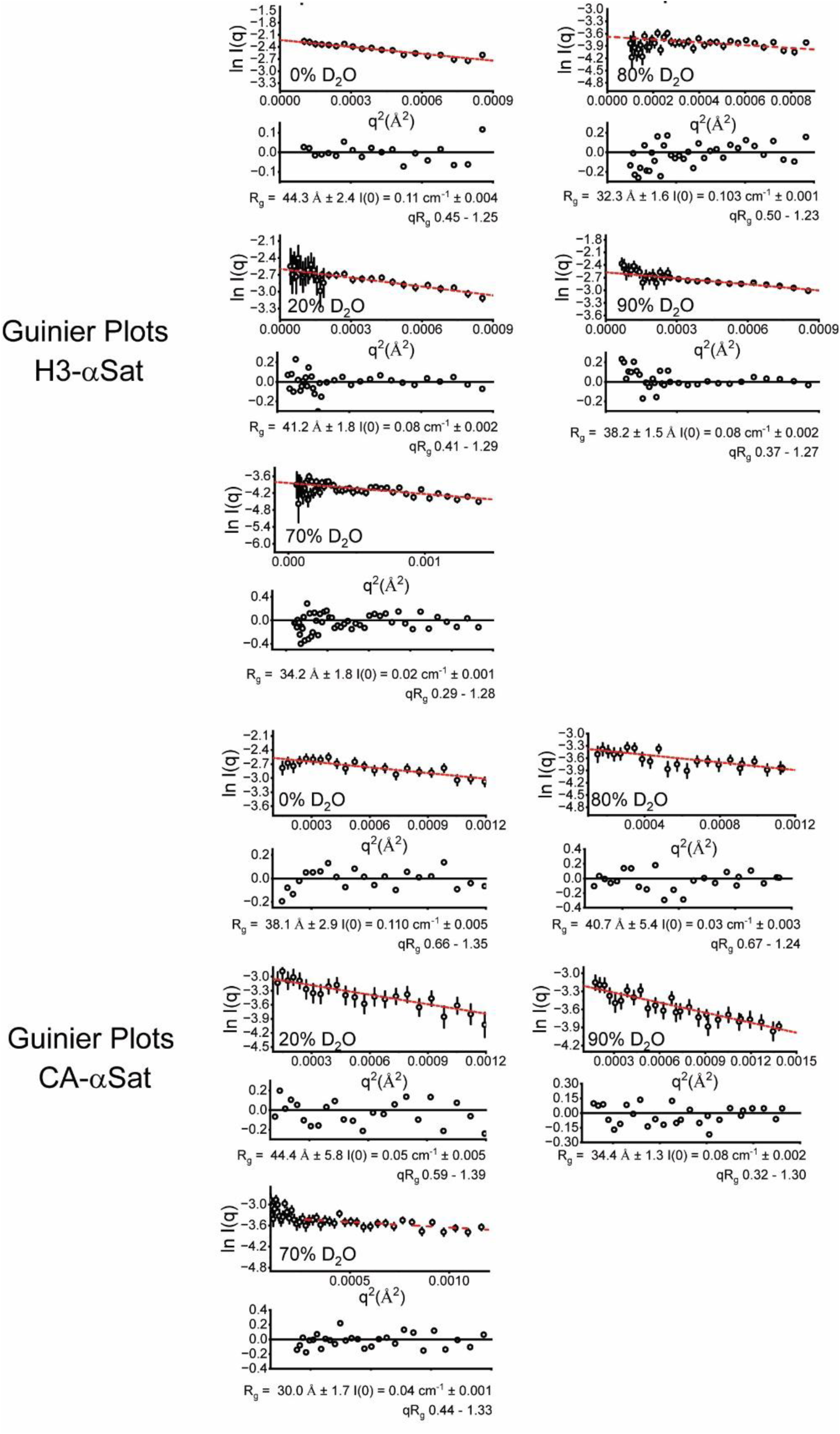
SANS Guinier Plots for H3-αSat and CENP-A-αSat. Guinier plot analyses (red lines) (ln (I) vs. *q*^2^) of SAXS data (black open circles) for H3-αSat and CENP-A-αSat NCPs, with residuals from the fitted lines shown below. Monodispersity is evidenced by linearity in the Guinier region of the scattering data and agreement of the I_0_ and R_g_ values determined with inverse Fourier transform analysis by the programs GNOM (Table 1). Guinier analyses were performed where qR_g_ ≤ 1.4.

**Supplemental Figure 5.**
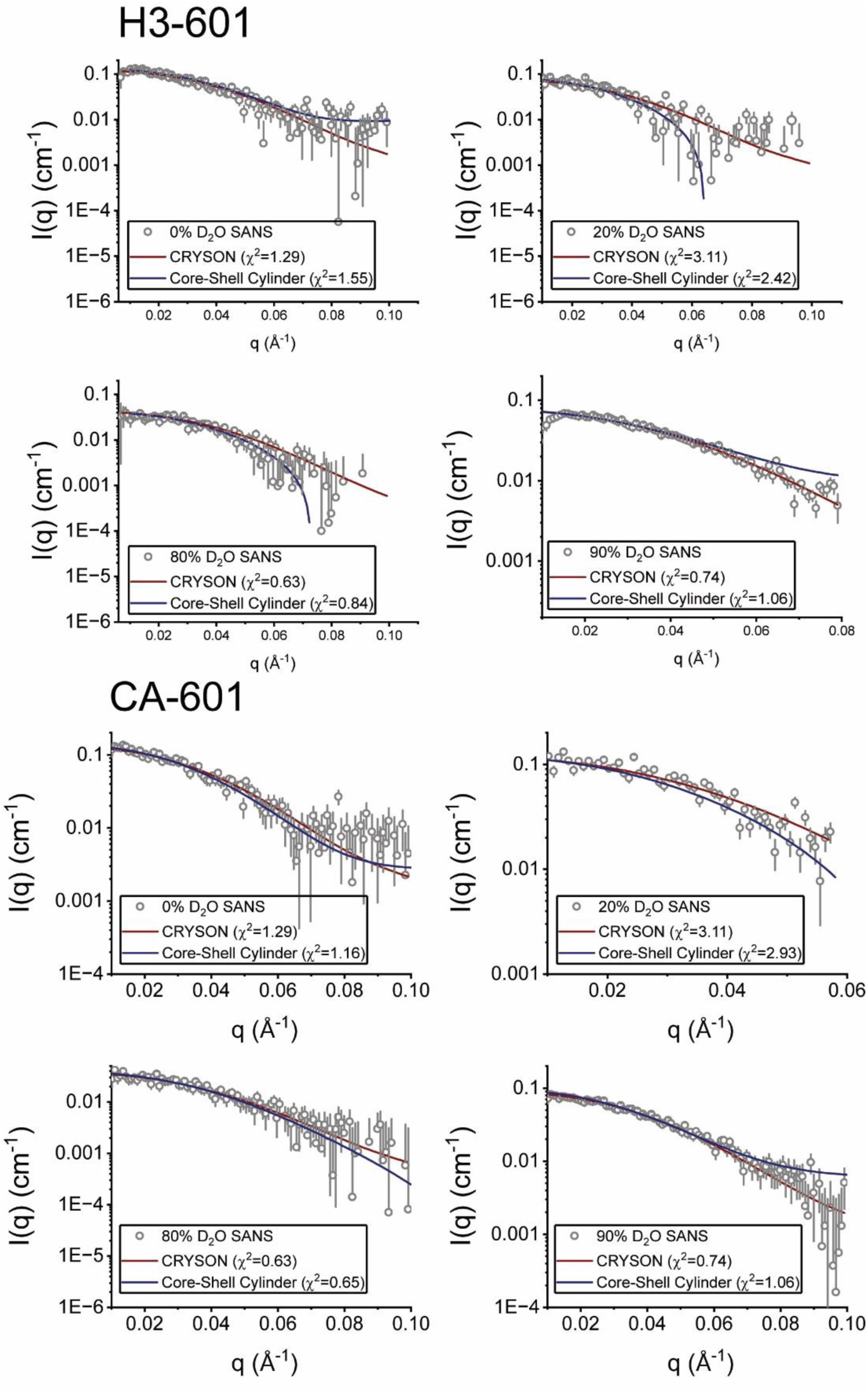
CRYSON and Core-Shell Ellipsoid Fitting of SANS Data for H3-601 and CENP-A-601. Shown are individual CRYSON fits (red lines) of the H3-601 or CENP-A-601 atomic models to experimental SANS data for both H3- and CENP-A 601 particles at each of four different D_2_O concentrations, with the χ^2^ for each respective fit provided in the graph legend. Fits were performed where q_max_ < 0.1 Å^−1^. Shown as blue lines are the global core-shell ellipsoid fits for SANS data for either the H3-601 or CENP-A-601 particles, as implemented in the program SASVIEW. See **Supplemental Table 4** for the structural parameters derived and individual and global χ^2^ associated with this fitting. Error bars on the scattering data (grey circles) represent the uncertainty associated with the intensity recorded.

**Supplemental Figure 6.**
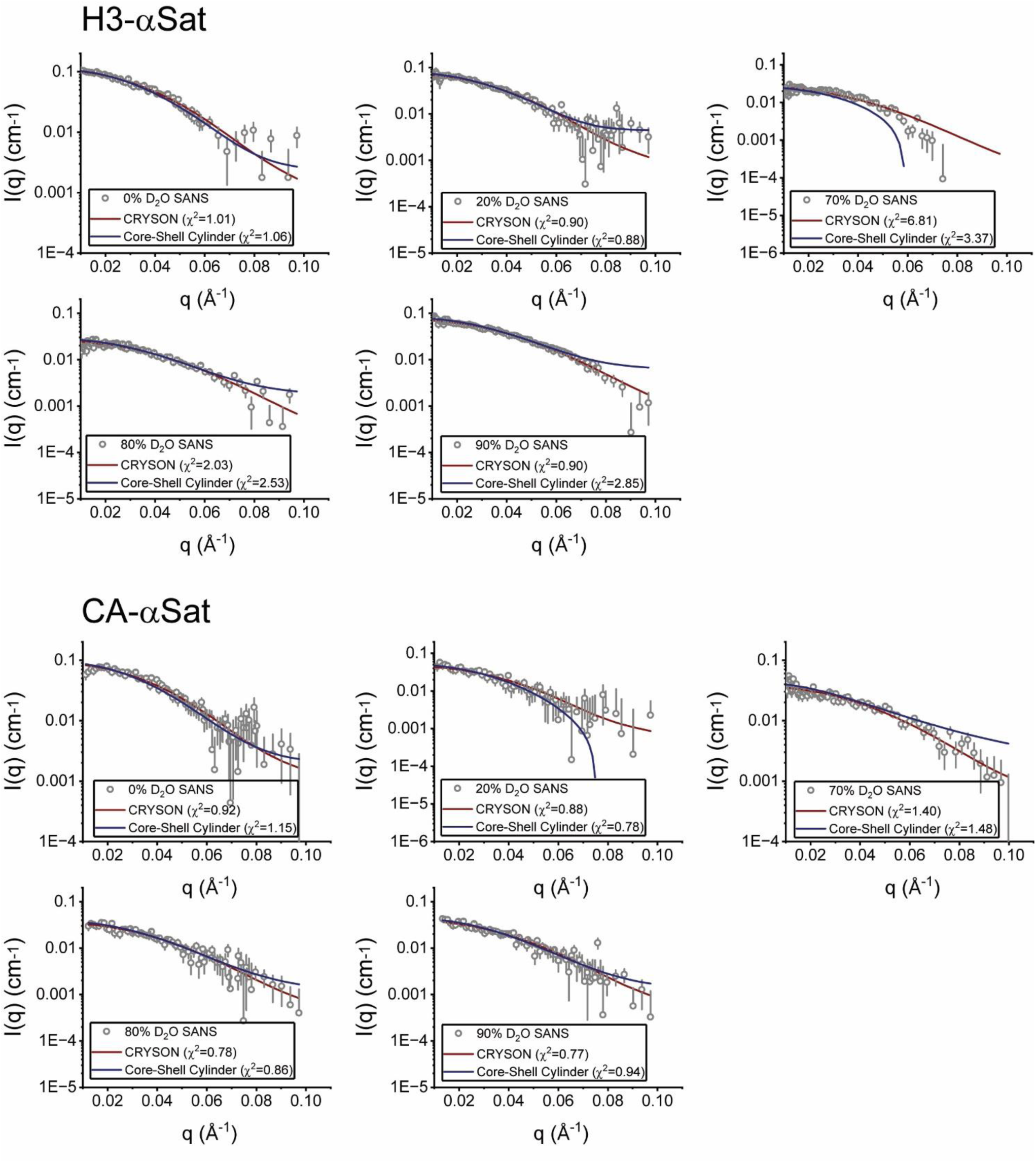
CRYSON and Core-Shell Ellipsoid Fitting of SANS Data for H3-αSat and CENP-A-αSat. Shown are individual CRYSON fits (red lines) of the H3-601 or CENP-A-601 atomic models to experimental SANS data for both H3- and CENP-A αSat particles at each of five different D_2_O concentrations, with the χ^2^ for each respective fit provided in the graph legend. Fits were performed where q_max_ < 0.1 Å^−1^. Shown as blue lines are the global core-shell ellipsoid fits for SANS data for either the H3-αSat or CENP-A-αSat particles, as implemented in the program SASVIEW. See **Supplemental Table 4** for the structural parameters derived and individual and global χ^2^ associated with this fitting. Error bars on the scattering data (grey circles) represent the uncertainty associated with the intensity recorded.

